# Mutations in Auxilin cause parkinsonism via impaired clathrin-mediated trafficking at the Golgi apparatus and synapse

**DOI:** 10.1101/830802

**Authors:** Dorien A. Roosen, Natalie Landeck, Luis Bonet-Ponce, Jillian Kluss, Melissa Conti, Nathan Smith, Sara Saez-Atienzar, Jinhui Ding, Aleksandra Beilina, Ravindran Kumaran, Alice Kaganovich, Johann du Hoffmann, Chad D. Williamson, David C. Gershlick, Luciana Sampieri, Christopher K. E. Bleck, Chengyu Liu, Juan S. Bonifacino, Yan Li, Patrick A. Lewis, Mark R. Cookson

## Abstract

Parkinson’s disease (PD) is a common neurodegenerative motor disorder characterized in part by neuropathological lesions in the nigrostriatal pathway. Loss of function mutations in Auxilin, the major neuronal clathrin uncoating protein, cause an aggressive form of juvenile onset PD. How mutations in Auxilin cause PD, is currently not understood. Here, we generated a novel mouse model carrying an endogenous pathogenic Auxilin mutation that phenocopies neurological features observed in patients, including motor impairments and seizures. Unbiased mapping of the Auxilin interactome identified synaptic and Golgi-resident clathrin adaptor proteins as novel interactors. Impaired clathrin-mediated trafficking in mutant Auxilin mice, both at the Golgi and the synapse, results in neuropathological lesions in the nigrostriatal pathway. Collectively, these results provide molecular mechanisms of PD pathogenesis in Auxilin mutation carriers, reinforcing a key role for clathrin-mediated trafficking in PD, and expand our understanding of the cellular function of Auxilin.

## Introduction

Parkinson’s disease (PD) is a multisystem neurodegenerative disorder characterized by a range of motor and non-motor neurological symptoms, including the clinically diagnostic triad of bradykinesia, rigidity and tremor (*1*). Although the neurochemical basis of PD has been understood for many decades, the underlying mechanisms driving the loss of neurons in the central nervous system of people with this disorder have remained, for the most part, obscure (*2*). This gap in our knowledge has contributed to the absence of disease modifying therapies for PD. The majority of PD cases are idiopathic in nature, however an estimated 5-10% of PD cases are inherited in a Mendelian fashion (*3*). Investigations into the function and dysfunction of genes identified in Mendelian PD have provided important insights into the etiology of Parkinson’s, highlighting protein aggregation, disruption of lysosomal biology, and mitochondrial quality control as key elements of the cellular events that lead to neurodegeneration (*2*, *3*).

Mutations in *DNAJC6/Auxilin* were first described in a series of young and juvenile onset patients with parkinsonism in 2012. The majority of these mutations are splice site mutations resulting in decreased Auxilin levels (*4*, *5*) or nonsense mutations resulting in a C-terminal truncation of Auxilin (*6*–*8*), which, along with the autosomal recessive (AR) mode of inheritance, support a loss of function mechanism driving disease in these cases, although one coding mutation has also been reported (R927G) (*5*). The clinical phenotype associated with mutations in *DNAJC6* is complex, with developmental delay, pyramidal symptoms and, in some cases, seizures in addition to a parkinsonian presentation (*4*–*8*).

Auxilin is involved in the uncoating and release of clathrin-coated vesicles (CCVs) through coordination of HSC70 chaperone activity and by interacting with a panel of adaptor proteins (APs) (*9*–*17*). Clathrin plays a central role in several cellular vesicle pathways, forming a cage-like structure surrounding vesicles, facilitating endocytosis and trafficking along the secretory pathway. Hence, *DNAJC6* mutations implicate clathrin-mediated trafficking in PD pathogenesis. In humans, Auxilin is expressed exclusively in the cells of the central nervous system, with its close paralog G-Cyclin Associated Kinase (GAK), being expressed ubiquitously (*18*, *19*). Notably, *GAK* has been implicated in idiopathic PD through genome wide association studies and protein interactome analyses – suggesting that both Auxilin and GAK have a conserved role in the survival of dopaminergic neurons in the human brain (*20*–*22*). The precise details of the mechanisms linking Auxilin to neurodegeneration have, however, yet to be elucidated.

Here, we set out to test the impact of coding variation in the *DNAJC6* gene causative for PD. We generated a new engineered rodent model to examine the organismal impact of the R927G mutation, in parallel with protein interactome analyses to shed light on the molecular basis for Auxilin dysfunction in PD. Our results provide the first evidence for neuronal dysfunction in a murine model for *DNAJC6* mutations and demonstrate a specialized role for Auxilin in the regulation and trafficking of CCVs derived from the *trans-Golgi* network (TGN) and in synaptic vesicle (SV) recycling. We show mechanistically that mutations in *DNAJC6*/Auxilin are loss of function through diminished interaction with its chaperone HSC70 as well as clathrin itself, even though interactions with clathrin adaptor proteins were retained. These data reveal important insights that expand the landscape of vesicle trafficking defects linked to PD, opening up novel pathways that are relevant for therapeutic development in this disorder, as well as illuminating the basic function and biology of Auxilin in CCV activity.

## Results

### Pathogenic R857G Auxilin allele is hypomorphic during early development

To analyze the impact of the human pathogenic R927G Auxilin mutation at the physiological level *in vivo*, we developed a homozygous knockin (KI) mouse model carrying the equivalent endogenous homozygous murine variant R857G (Figure 1A, B; Figure 1S). Analysis of Auxilin RNA expression in WT mice confirmed expression in dopaminergic neurons in the nigrostriatal pathway, a major area impacted during PD pathogenesis (Figure 1C).

**Figure 1:**
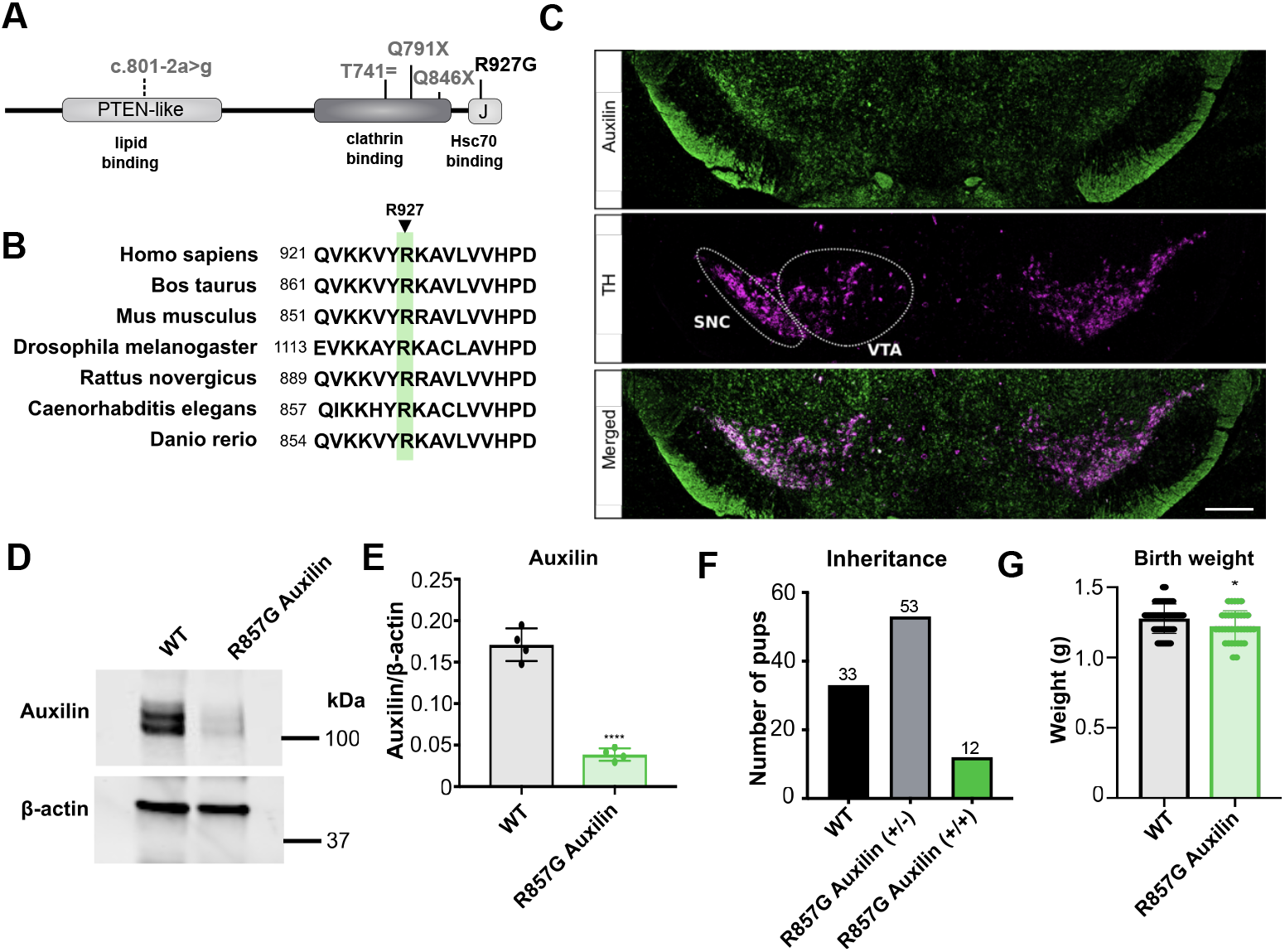
Novel PD mouse model with endogenous R857G Auxilin mutation displays hypomorphism during early development. A Overview of Auxilin functional domains and pathogenic mutations. B The human R927 Auxilin residue is conserved across species and is equivalent to the murine R857 residue. C RNAscope of midbrain slices of 2 month old WT mice indicating Auxilin mRNA (green) and the dopaminergic neuronal marker TH mRNA (pink). Scale bar indicates 800 μm. SNC substantial migration pars compacta, VTA ventral tegmental area. E WB of primary neurons of WT and R857G Auxilin mice. F Quantification of normalized Auxilin levels of n=4 independent cultures. Welch’s t-tests were performed, **** indicates p-value < 0.0001. G Survival bias of the offspring of heterozygous R857G Auxilin mating pairs at 3 weeks old. Chi-square test was performed with n = 10 litters, with resulting p-value < 0.01. H Body weight of newborn mice, Welch’s t-test was performed of n=5 and n=6 litters for WT and R857G Auxilin mice, respectively, * indicates p-value < 0.05.

Analysis of Auxilin protein levels in primary neurons derived from newborn (p0) R857G Auxilin mice revealed lower Auxilin protein expression compared to WT controls (Figure 1D, E). Similarly, brain lysates of p0 as well as p2 R857G Auxilin mice had lower Auxilin protein levels than WT animals, with some variation between animals (Figure S2 A, B, D, E). However, no differences in Auxilin protein levels were observed in the brain of p6 or 3 week old mice (Figure S2 G, H, J, K), indicating an age-dependent upregulation of Auxilin during early development of R857G Auxilin mice. GAK, the ubiquitously expressed paralogue of Auxilin, has previously been found to be upregulated in the brain of Auxilin mutation carriers as well as Auxilin KO mice, likely to compensate for loss Auxilin function (*8*, *23*). Hence, we analyzed GAK protein levels in the brain of R857G Auxilin mice and observed a modest, transient upregulation of GAK protein in the brain of R857G Auxilin mice during early development (Figure S2 D, F). Similar to conventional Auxilin KO mice (*23*), R857G Auxilin mice displayed increased perinatal mortality, as indicated by a deviation from the expected 1:2:1 inheritance from heterozygous breeding pairs 3 weeks after birth (Figure 1F). Birth weights of R857G Auxilin mice were also lower than their wild type counterparts (Figure 1G). These results suggest that R857G Auxilin is a loss of function allele *in vivo* but that upregulation of the mutant variant occurs, presumably to allow survival of mice carrying homozygous alleles.

### Neurological phenotypes in R857G Auxilin mice phenocopy clinical features seen in patients

To assess neurological phenotypes in R857G Auxilin mice, we used a battery of behavioral tests on a longitudinal cohort of 8 mice per genotype at 6, 12 and 18 (Figures S4, 2 and S5, respectively) months of age. In contrast with the slightly decreased weight at birth, mutant Auxilin mice exhibited no weight differences at these ages (data not shown). R857G Auxilin mice displayed balance impairments, as indicated by an increased tendency to fall from an elevated beam during the beam walk test compared to WT mice (Figure 2A; Figure S4A, S5A). The pole test revealed progressive bradykinesia and decreased agility from 12 months onward in R857G Auxilin mice, as shown by an increased time to turn and to descend from a vertical wooden pole (Figure 2B, C; Figure S4 B,C; S5 B, C). However, R857G Auxilin mice outperformed WT mice during the rotarod test, suggesting that gross motor function is overall intact in these animals (Figure 2D; Figure S4D, S5D). Analysis of the amplitude of movement over a broad-frequency range in a startle chamber revealed a decreased amplitude of movement in 12-month-old R857G Auxilin mice (Figure 2 E-G), which along with the pole test indicates bradykinesia. A subset of R857G Auxilin mice were observed to suffer seizures during cage changes (Video S1). Seizures were characterized by a freezing phenotype, followed by drooling and muscular twitching (Video S1). R857G Auxilin mice of up to 18 months of age were not observed to suffer from anxiety, memory, decreased forelimb strength or locomotor phenotypes, as assessed by the Y-maze, elevated plus maze, grip strength and open field tests (results shown for 12 month old mice, Figure S3). Taken together, these behavioral tests show R857G Auxilin mice develop neurological phenotypes that phenocopy clinical features seen in patients harboring mutations in *DNAJC6*, including typical parkinsonian motor impairments, such as progressive bradykinesia and gait disturbances, as well as seizures. However, as many behaviors remain unaffected, the impairments are specific to a subset of motor functions.

**Figure 2:**
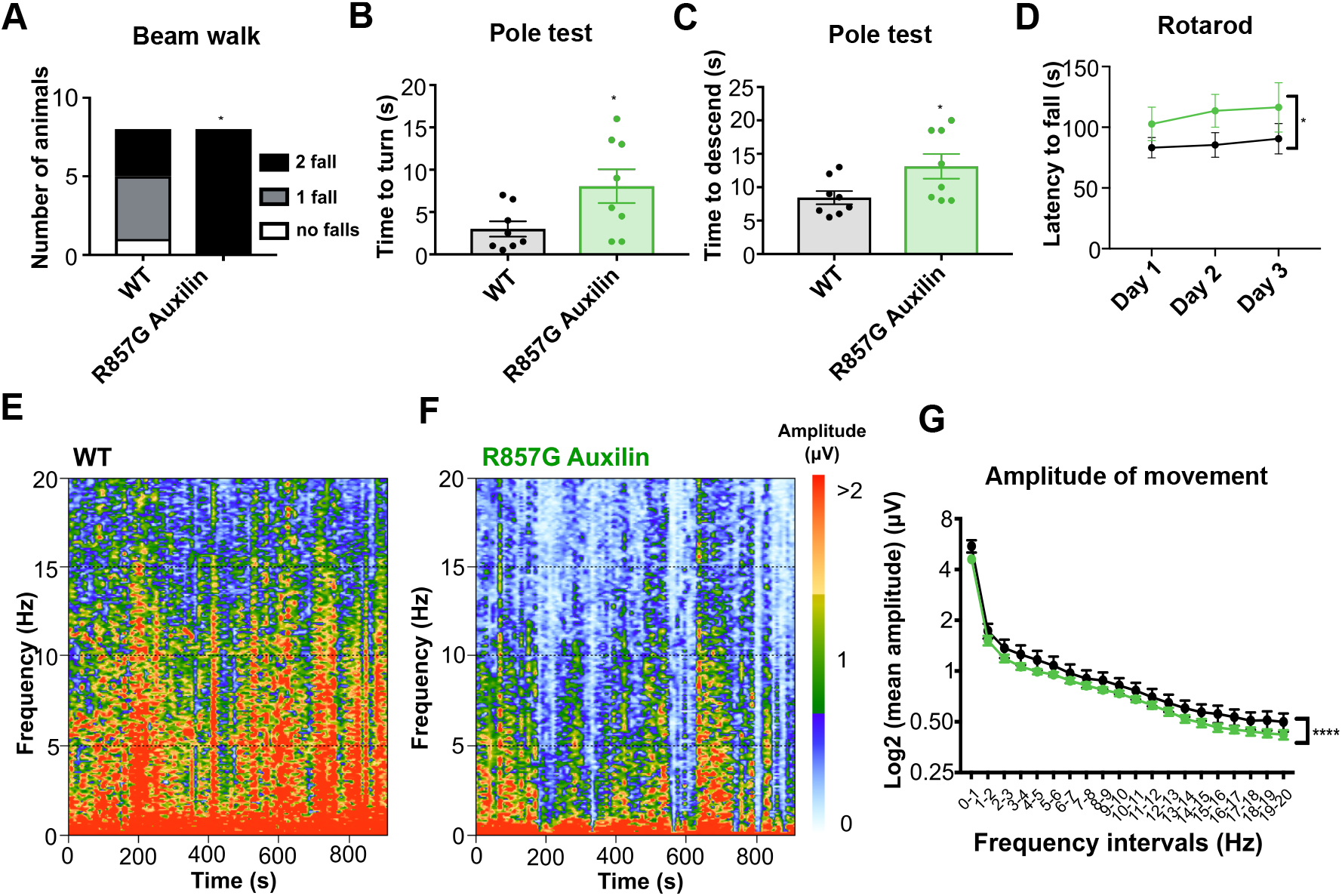
Motor impairments in 1 year old R857G Auxilin mice phenocopy clinical features observed in patients. A Beam walk analysis indicating the number of falls when traversing a 6 mm beam during 2 trials. Chi-square tests were performed, * indicates p < 0.05. B, C Performance during the pole test as analyzed by time to turn and time to descent. Welch’s t-tests were performed with * p < 0.05. D Latency to fall during the rotarod test during 3 consecutive days. Two-way ANOVA was performed with * p < 0.05. E, F Analysis of the amplitude of movement in a startle chamber. Representative spectrographs of WT and R857G Auxilin mice are shown, displaying the amplitude of movement (indicated by color) during 15 minutes over a frequency range of 0-20 Hz. G Two-way ANOVA was performed, with **** p < 0.0001.

### Auxilin interacts with clathrin adaptor proteins at the TGN and at the synapse

To gain further insight into the molecular mechanisms contributing into Auxilin-associated PD pathogenesis, the interactome of Auxilin was mapped in an unbiased fashion by combining GFP-nanotrap affinity purification with SILAC-based proteomics (Figure S6A). Experiments were performed in triplicate (figure S6B) and bio-informatic filtering of protein interactors identified across all replicates resulted in a total of 32 top candidate Auxilin interactors (Figure 3A). Among the identified interactors were Auxilin itself and previously reported interactors clathrin heavy chain (CHC) and the plasma-membrane resident clathrin adaptor protein AP2 subunit α2 (AP2α2) (*24*), indicating that the experiment was successful in recovering authentic Auxilin interactors (Figure 3A). HSC70, the chaperone of Auxilin required for its role in CCV uncoating, is another *bona fide* Auxilin interactor that was identified by mass spectrometry (-log2(SILAC ratio) of 3.028 and −log10(FDR-corrected P-value) of 2.30). However, HSC70 was excluded from the final list of Auxilin interactor top candidates by bioinformatic filtering, due to its high abundance (>95%) in repositories of common contaminants of affinity purification-mass spectrometry experiments(*25*).

**Figure 3:**
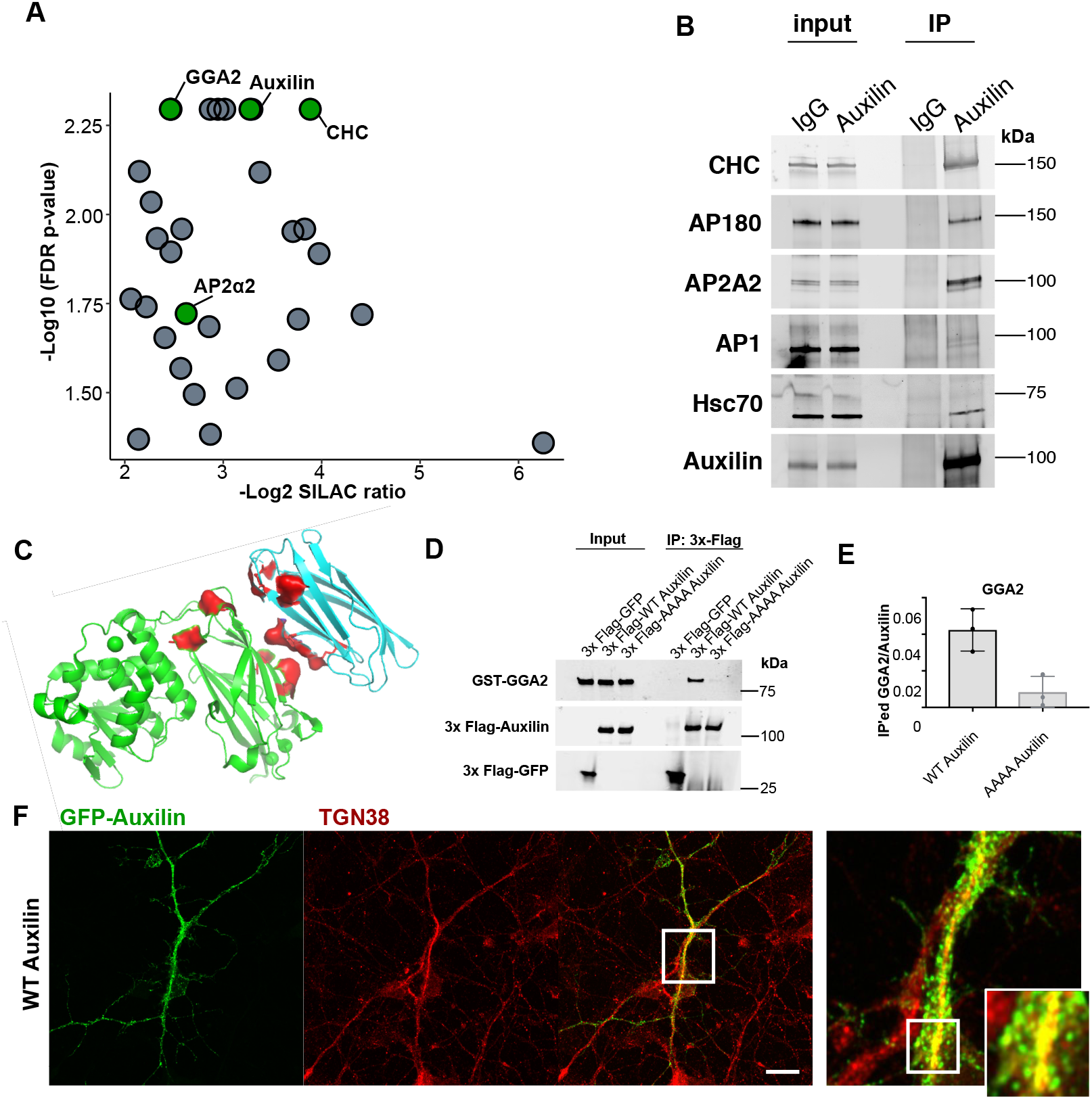
Auxilin interacts with a specific subset of synaptic and Golgi-resident clathrin adaptor proteins. A Scatter plot of top candidates Auxilin interactors. B Immunoprecipitation of endogenous Auxilin from mouse brain, probed for interaction with CHC, HSC70, AP180, AP2A2 and AP1. C Structural model of bovine Auxilin (green) with GGA2 (cyan). Putative interacting motifs are indicated in red (F and L from FIPL motifs in Auxilin, AR, K, RR from conserved surface motifs in GGA2 (*28*). D Recombinant 3xFlag-GFP, 3xFlag-WT Auxilin or 3xFlag-AAAA Auxilin were incubated with recombinant GST-GGA2, followed by 3xFlag co-immunoprecipitation and WB analysis. Image is representative of n = 3 technical replicates. E Quantification of GGA2 co-purified with 3xFlag-Auxilin, normalized to bait. Unpaired t-test was performed and * is p-value < 0.05. F Representative confocal image of primary murine neurons transiently expressing GFP-Auxilin (green) and co-stained for the endogenous TGN marker TGN38 (red). Scale bar = 20 μm.

We sought to validate Auxilin interactors at the endogenous level by performing coimmunoprecipitation of Auxilin from mouse brain. We confirmed that Auxilin interacts with its chaperone HSC70 and clathrin, as well as the plasma membrane-resident adaptor protein AP2 but to a much lower extend with the Golgi-resident adaptor protein AP1 (Figure 3B). In addition, we queried the interaction of Auxilin with neuron-specific clathrin adaptor proteins, that might not have been recovered in the proteomics screen in HEK cells. This identified AP180 as a novel interactor of Auxilin (Figure 3B), as its expression is restricted to neurons, required for the recycling of synaptic vesicles.

In contrast with its ubiquitously expressed homologue GAK, Auxilin has not previously been reported to interact with Golgi-resident proteins. Remarkably however, gene ontology analysis of the Auxilin interactome indicated an enrichment both plasma membrane and Golgi apparatus-associated cellular components (Figure S6 C, D). Indeed, the Golgi-resident clathrin adaptor protein GGA2 was identified as a novel Auxilin interactor candidate (Figure 3A).

To further compare and validate the interaction of Auxilin and its homologue GAK with multiple clathrin adaptor proteins, co-IPs were performed from HEK293FT cells transiently expressing GFP-Auxilin or GFP-GAK and WBs were probed for all endogenous adaptor proteins with a reported role in clathrin-mediated trafficking (AP1-3 and GGA1-3) (Figure S7A-F). We confirmed the interaction of Auxilin and GAK with AP2, and of GAK but not Auxilin with AP1, as previously reported (Figure S7 A, C, D) (*16*, *17*, *24*, *26*). No interaction was observed between Auxilin or GAK with AP3 (Figure S7 A). In addition, co-IP confirmed interaction between Auxilin and GGA2 and we also observed a weak interaction with GGA3, but not GGA1 (Figure S7 B, E, F). GAK was not found to interact with any of the GGA proteins, as previously reported (Figure S7 B, E, F) (*16*). These data indicate that Auxilin and GAK both interact with the plasma membrane-resident AP2 but display differential binding with Golgi-resident clathrin adaptor proteins. Altogether, these findings indicate for the first time a direct role for Auxilin in the uncoating of Golgi-derived CCVs, via interaction with the Golgi-resident adaptor protein GGA2. To test whether these interaction data are supported by spatial proximity within the cell using an alternative method, we transiently transfected GFP-Auxilin and examined co-localization with the TGN marker TGN38 (Figure 3 F). The co-localization of Auxilin with this marker further supports a physiologically relevant function for Auxilin at the Golgi apparatus.

### Identification of binding motifs required for differential interaction of Auxilin and GAK with Golgi-resident clathrin adaptor proteins

The γ-subunit of AP1 and the GGAs are γ-ear containing proteins that have previously been found to interact with accessory proteins through a conserved consensus motif ψG(P/D/E)(ψ/L/M) (*27*). GAK interacts with AP1-γ via two sequences fitting this motif, namely FGPL and FGEF. These sequences are not conserved in Auxilin, which could explain its inability to interact with AP1 (Figure S8A). Further analysis of the Auxilin amino acid sequence revealed that Auxilin contains two sequences closely resembling the ψG(P/D/E)(ψ/L/M) consensus motif, namely FIPL (Figure S8B). Structural modelling of Auxilin and GGA2 shows the presence of the FIPL motifs on the surface of the tertiary structure and can be modelled to fit in close proximity with the basic surface residues of GGA2 required for recruitment of accessory proteins (Figure 3C). *In vitro* protein binding assay of recombinant 3x-Flag-Auxilin with GST-GGA2 showed a direct interaction between both proteins, making GGA2 a *bona fide* interactor of Auxilin. Mutagenesis of the putative GGA-binding motifs FIPL to AIPA (Figure S8C) markedly reduced interaction of Auxilin with GGA2, indicating that these motifs are indeed required for the interaction (Figure 3 D, E). The FIPL motifs are not conserved in the GAK amino acid sequence, which may explain the inability of GAK to interact with GGA2 (Figure S8B).

### Pathogenic Auxilin mutations impair interaction with clathrin

Next, we set out to address the impact of PD-associated mutations on the interactome of Auxilin. Endogenous Auxilin was co-immunoprecipitated from WT and R857G mutant mouse brain. The R857G mutation, equivalent to the R927G mutation in human Auxilin, resides within the J-domain of Auxilin, required for the interaction with its chaperone HSC70 (Figure S11A). Indeed, we found that the R857G mutation reduced the interaction of Auxilin with HSC70 (Figure 4 A, E). Furthermore, interaction of mutant Auxilin with clathrin was also impaired (Figure 4 A, D), whereas interaction with clathrin adaptor proteins AP2 and AP180 remained unaltered (Figure 4 A, F, G) To further address the impact of additional pathogenic mutations of human Auxilin on its interactors, WT GFP-Auxilin or PD-associated coding mutants (Q791X, Q846X, R927G) were transiently expressed in cells and WB was performed on immunoprecipitates. Interaction of the pathogenic Auxilin mutants with clathrin adaptor proteins AP2, GGA2 and GGA3 remained unaltered (Figure S7G, J-L). Truncating mutations Q791X and Q846X, which lack some of the clathrin binding boxes (Figure S11A), resulted in a marked decrease of clathrin interaction (Figure S7H, I). Like the R857G mutation in mouse Auxilin, the point mutation R927G also impaired interaction of Auxilin with clathrin. (Figure S7H, I). The R927G mutation resides within the J-domain of Auxilin (Figure S11A) (*29*, *30*); however, structural modelling of the J-domain with the clathrin coat shows that R927 is not in close proximity with the clathrin coat (Figure 4 B, C). It is of note that the J-domain is composed of highly conserved alpha-helical structures, relying on the formation of hydrogen bonds between negatively and positively charged amino acids. Substitution of the positively charged R927 side chain to the uncharged and smaller G927 would therefore be likely to disrupt the tightly packed helical structure, resulting in a decreased ability of correct positioning within the assembled clathrin coat (Figure 4C). Even though the clathrin binding boxes appear the main determinants for clathrin interaction (Figure S11A) (*17*, *24*), additional low affinity interaction with the J-domain may further contribute to Auxilin interaction with the clathrin lattice. Despite the reduction of clathrin binding by pathogenic Auxilin mutants, transient expression of WT as well as mutant (Q791X, Q846X, R927G) GFP-Auxilin in primary neurons showed colocalization with endogenous CCVs (Figure S9). This indicates that pathogenic Auxilin mutants can still be recruited correctly to CCVs, probably via interaction with lipids and clathrin adaptor proteins, but may be inefficient in the CCV uncoating reaction, as Auxilin interaction with both HSC70 and clathrin is crucial for the correct positioning of HSC70 chaperone to mediate efficient clathrin uncoating.

**Figure 4:**
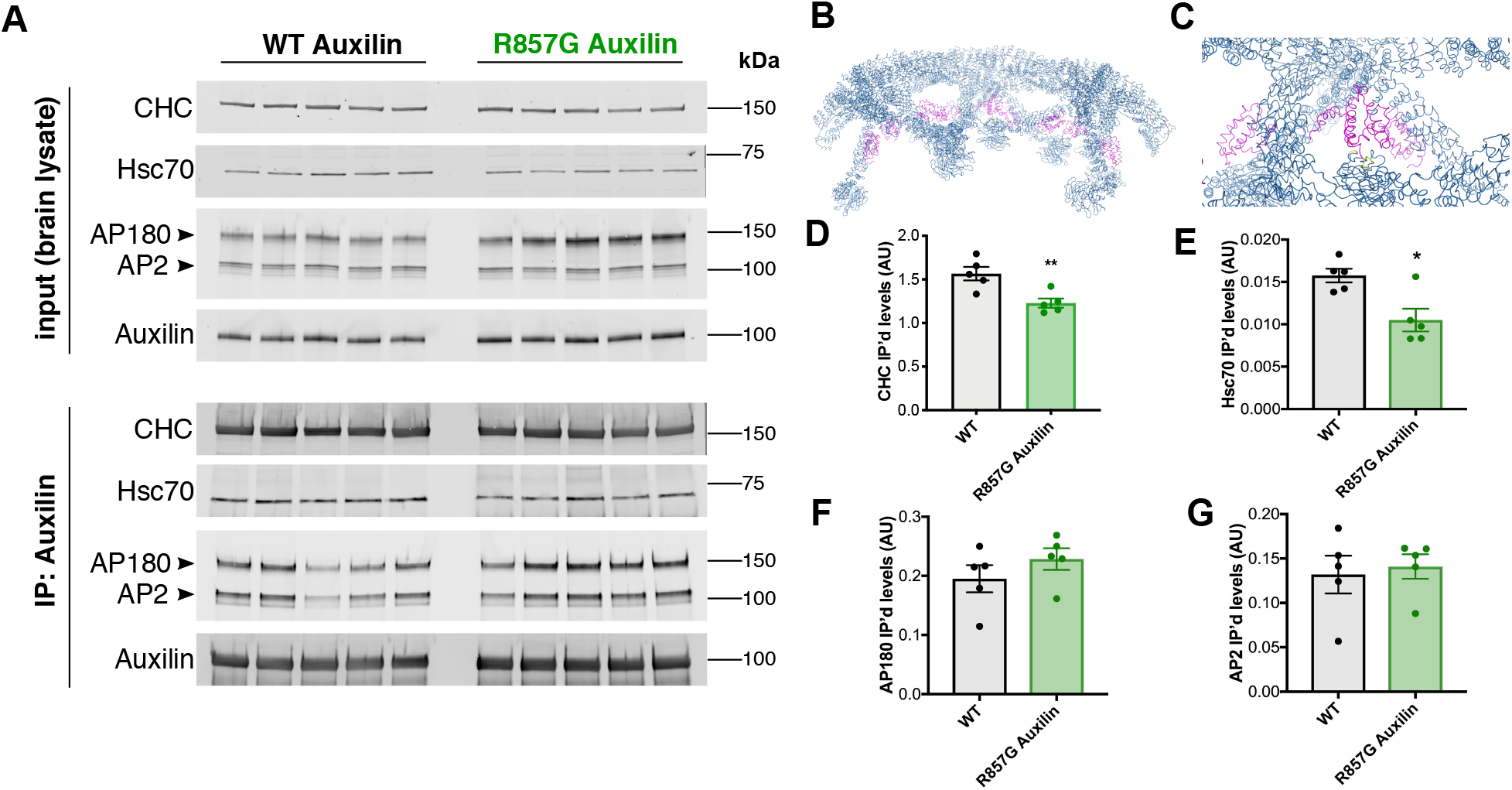
Impact of mutations on the interactome of Auxilin. A Immunoprecipitation of endogenous Auxilin from brain of WT or R857G Auxilin mice. Immunoprecipitates were probed for interaction of Auxilin with endogenous CHC, HSC70, AP2 and AP180. B, C J-domain of bovine Auxilin (purple) bound to the assembled clathrin coat (dark blue) (PDB file derived from (*11*). Auxilin residues residing at the interface with the clathrin coat (G825, P908, L909 and Y910, equivalent to human G885, P968, L969 and Y 970) are indicated in yellow. PD mutation (bovine R867, equivalent to human R927) is indicated in cyan. D-G Quantification of CHC, HSC70, AP180 and AP2, respectively, coimmunoprecipitated with Auxilin from WT and R857G Auxilin brain of n= 5 animals. Unpaired t-test was performed and * is p-value < 0.05, ** is p-value < 0.01.

### Impaired clathrin uncoating and synaptic vesicle recycling R857G Auxilin mice

PD is typically characterized by loss of dopaminergic (DA) neurons in the nigrostriatal pathway. Immunohistochemical analysis of tyrosine hydroxylase, a DA marker, did not reveal alterations in the *substantia nigra* or striatum of R857G Auxilin mice up to 12 months of age (Figure S10). The observed parkinsonian manifestations in R857G Auxilin mice are, therefore, not likely to be the result of gross dopaminergic neurodegeneration.

Interaction data have indicated that mutant Auxilin results in the impairment of interaction with HSC70 and clathrin. Therefore, we hypothesized that the uncoating reaction might stall, resulting in a decreased turnover of CCVs and hence to neuronal dysfunction. As Auxilin was found to interact with the synaptic clathrin adaptor proteins AP2 and AP180, and Golgi-resident adaptor protein GGA2, we sought to investigate CCV uncoating both at the synapse and the Golgi apparatus.

To investigate the molecular impact of the R857G Auxilin mutation on clathrin structures and synaptic function, the ultra-structural morphology of nerve terminals was studied in the mutant mouse brain using electron microscopy. Since neurological defects were observed in R857G Auxilin mice as early as 6 months, this analysis was carried out in the dorsal striatum of 6 month old mice. Whereas the synaptic area of neurons in striatal section was unaltered, the number of synaptic vesicles (SVs) at the presynaptic compartment was found to be decreased in R857G Auxilin mice (Figure 5A-D). The overall decrease in number of SVs suggests that synaptic recycling is impaired in mutant Auxilin mice due to inefficient uncoating of CCVs, as has previously been observed in mice with disrupted clathrin-associated endocytic proteins including Auxilin, Synaptojanin 1 and Endophilin 1 (*23*, *31*–*34*). To show the impairment of clathrin uncoating in R857G Auxilin brain, CCVs were isolated from WT and mutant mouse brain. We observed a striking increase of CCVs isolated from mutant Auxilin brain, as indicated by a drastic increase of clathrin in the pure CCV fraction (Figure 5F, G). Furthermore, AP180, a SV marker, was also increased in the CCV fraction, indicating that the recycling of SVs is indeed impaired (Figure 5 F, G). Auxilin levels remained unaltered (Figure 5 G), however the ratio of Auxilin over clathrin was decreased (Figure 5 H), in line with the impaired interaction of Auxilin with clathrin. To further analyze clathrin-mediated endocytosis in the context of synaptic recycling, we employed the well-established FM 1-43 dye endocytosis assay(*35*). Primary hippocampal neurons derived from WT and R857G Auxilin mice were cultured, stimulated and incubated with FM1-43 dye to label endocytosed synaptic vesicles. Quantification of FM 1-43 fluorescence intensity showed an accumulation of FM 1-43 in R857G mutant Auxilin neurons compared to WT, indicating the impaired recycling of synaptic vesicles, with subsequent intracellular trapping of intracellular FM 1-43 (Figure 5 I - L).

**Figure 5:**
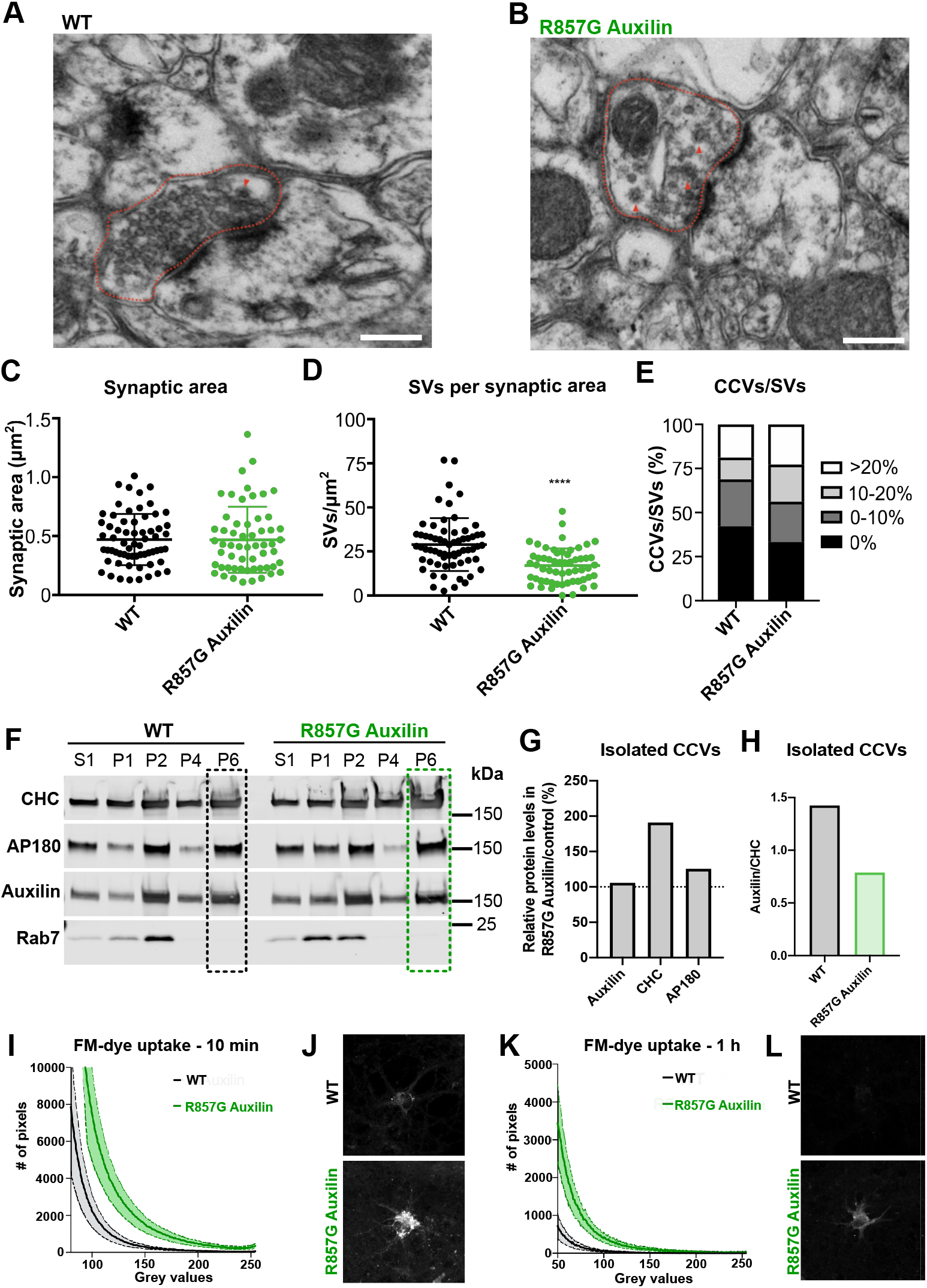
Impaired presynaptic vesicular recycling. A, B Representative EM images of synaptic terminals in the striatum of WT and R857G Auxilin mice, respectively. Presynaptic areas are marked by red dotted lines and CCVs by red arrows. Scale bar indicates 300 nm. C Quantification of the presynaptic area. D Quantification of the number of SVs per presynaptic area. N = 64 and n = 57 synapses were analyzed for WT and R857G Auxilin mice, respectively and Welch’s t-test was performed, with **** indicating a p-value < 0.0001. E Quantification of the percentage of CCVs in the pre-synaptic area over the number of SVs. D Quantification of the number of synaptic vesicles per pre-synaptic area. F Isolation of CCVs from the brain of WT and R857G Auxilin mice, with P6 indicating the clathrin-pure fraction. Clathrin fraction was probed for CHC, AP180, Auxilin and Rab7 as a negative control. N = 20 mice per genotype. G Quantification of the percentage of Auxilin, CHC and AP180 protein levels in the clathrin-pure fraction of CCVs isolated from R857G Auxilin brain compared to WT control. H Quantification of Auxilin in the CCV-purified fraction. F, K Quantification of FM-dye uptake in primary neurons derived from WT and R857G Auxilin mice after 10 and 60 min, respectively. J, K Representative confocal images of FM 1-43 labeling of WT or R857G Auxilin primary neurons.

#### Impaired uncoating of Golgi-derived CCVs and dysmorphic Golgi morphology in R857G Auxilin mice

In addition to an increased number of CCVs in the nerve terminals of R857G mice, we also observed a significant increase of coated structures around the Golgi apparatus (Figure 6 A-C). Further EM analysis of Golgi apparatus in the dorsal striatum of R857G Auxilin mice revealed dystrophic morphological alterations of the Golgi apparatus. Golgi stacks in R857G Auxilin mice often appeared more swollen as compared to WT controls (Figure 6 A, B, F). To be able to differentiate between *cis*, medial and *trans*-Golgi stacks, murine primary neurons were stained for endogenous Golgi markers and analyzed with enhanced-resolution microscopy. Swollen Golgi morphology would result in an increased surface area, with subsequent decrease of colocalization between neighboring Golgi stacks. Consistent with the hypothesis of Golgi swelling, decreased co-localization between cis and medial Golgi stacks (GM130 and GLG1, respectively) and medial and *trans*-Golgi stacks (GLG1 and TGN38, respectively) was observed in primary neurons derived from R857G Auxilin mice compared to WT (Figure 6 D, E, G, H). Taken together, our findings indicate that mutant Auxilin impairs uncoating of CCVs both at the synapse and the Golgi apparatus.

**Figure 6:**
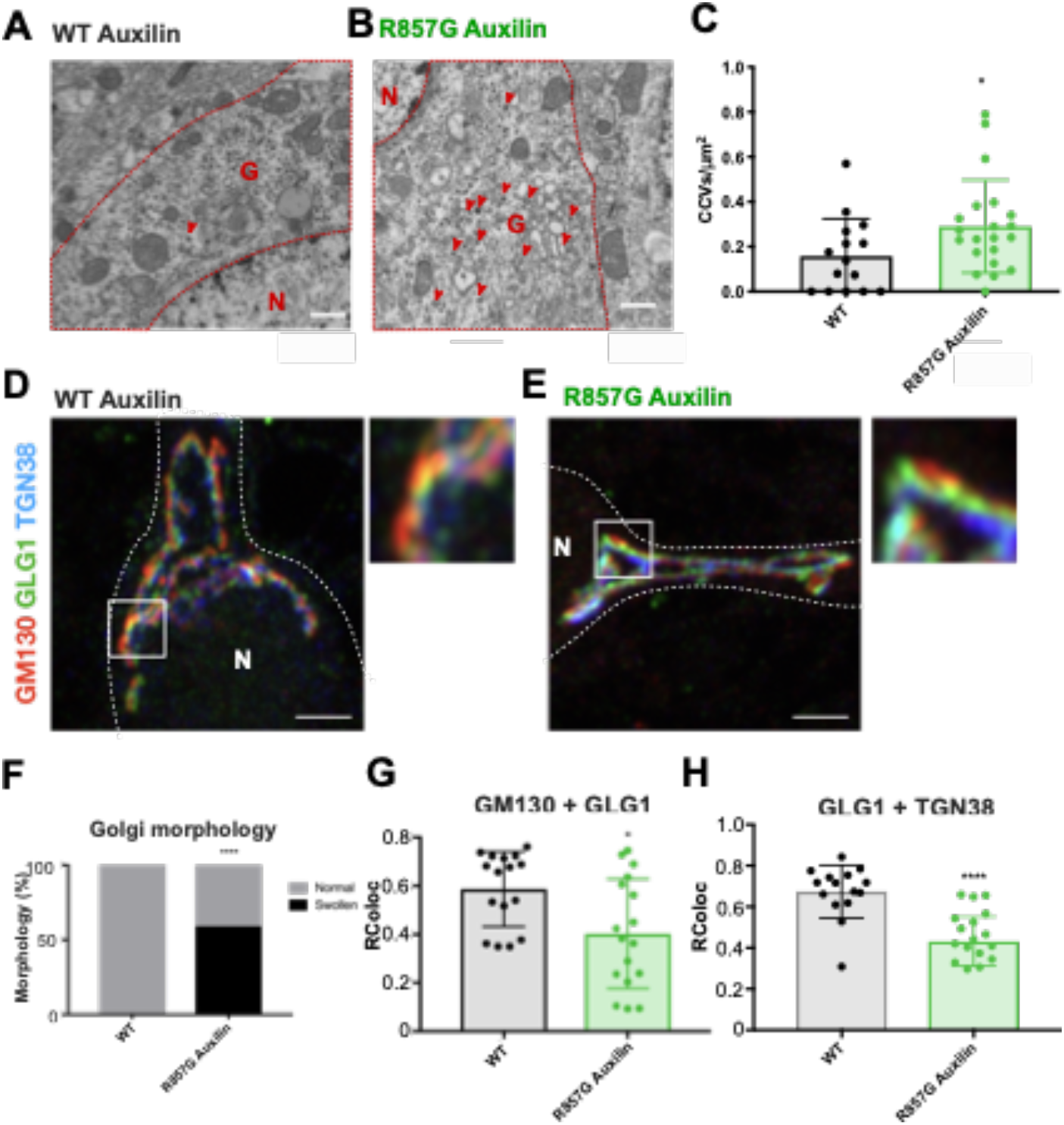
Accumulation of coated structures and dystrophic morphological alterations of the Golgi apparatus. A, B Representative EM images of brain slices of the striatum of WT and R857G Auxilin mice. Scale bar = 600 nm. Cytoplasm is marked by red dotted lines. N = nucleus, G = Golgi. Red arrows indicate CCVs. C Quantification of the number of CCVs per cytoplasmic area. Welch’s t-test was performed, n= 15 and n = 21 cells were analyzed for WT and R857G Auxilin mice, respectively. D, E Representative confocal images with Airyscan detection of WT and R857G Auxilin primary neurons, stained for endogenous GM130 (red), GLG1 (green) and TGN38 (blue). Scale bars = 2 μm. N is nucleus, neuronal cell body is indicated in white dotted line. F Quantification of the observed Golgi morphologies of n = 10 and n = 22 WT and R857G Auxilin striatal cells in A and B. Binomial tests were performed, **** indicates p-value < 0.0001. G, H Quantification of the co-localization of GLG1+GM130 and GLG1+TGN38 of n=16 WT and n=17 R857G Auxillin primary neurons. Unpaired t-tests were performed, * indicates p-value < 0.05, **** indicates p-value < 0.0001.

### PD-like neuropathology in the striatum of R857G Auxilin mice

Efficient uncoating of Golgi-derived CCVs is a key event in the correct delivery of lysosomal proteins. Therefore, we hypothesized that impaired clathrin-mediated trafficking in R857G Auxilin mice may result in the accumulation of endolysosomal cargo in neurons. Remarkably, EM analysis revealed the presence of large intracellular lipid/proteinaceous aggregates reminiscent of lipofuscin in the dorsal striatum of R857G Auxilin mice (Figure 7 A). Progressive accumulation of lipofuscin has previously been observed in familial PD cases, pointing to impairments of the lysosomal system (*2*, *36*). To further assess the accumulation of cargo, midbrain sections were stained using BODIPY for neutral lipids in dopaminergic neurons in the *substantia nigra.* Neurons in R857G Auxilin mice contained less lipid droplets, but the size of each droplet was significantly increased, resulting in an overall increase of total lipid content per dopaminergic neuron in the SN of R857G Auxilin mice (Figure 7 B-F).

**Figure 7:**
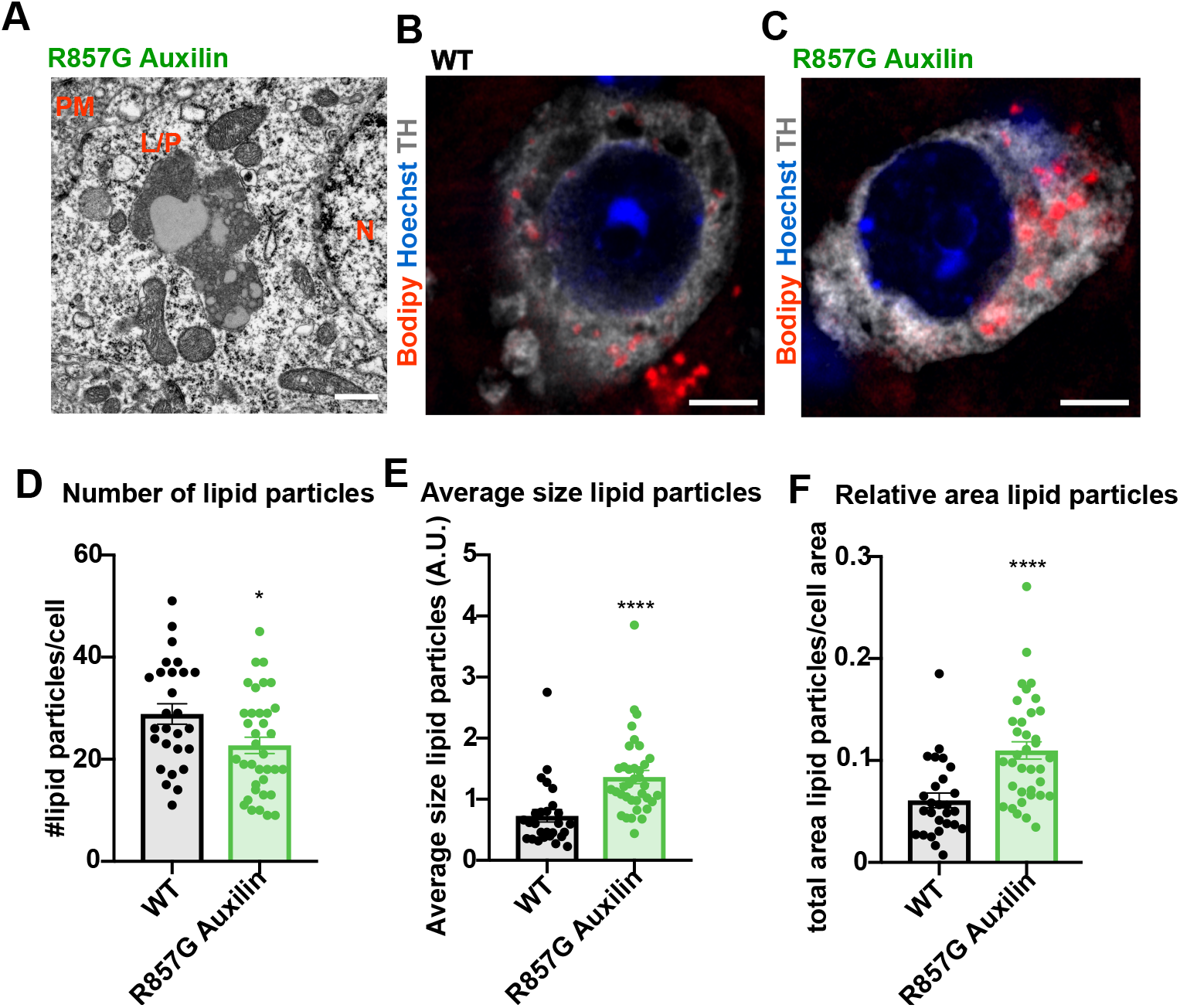
Accumulation of lipofuscin and lipids in neurons of R857G Auxilin mice. A EM analysis of striatal sections of a R857G Auxilin mouse. PM indicates plasma membrane, L/P lipid protein aggregate, N nucleus. Scale bar indicates 300 nm. B, C BODIPY staining and confocal imaging with Airyscan detection of DA neurons in sections of the SN of WT and R857G Auxilin mice. Scale bar indicates 5 μm. D, E, F Quantification of number of particles, particle size and lipid content per DA cell in the SN, respectively. 27 and 36 cells of n = 3 and n = 4 WT and R857G Auxilin mice, respectively, were analyzed. Welch’s t-test was performed, * indicates p-value <0.05, **** indicates p-value <0.0001.

## Discussion

Over the past two decades, a number of molecular pathways have been linked to the pathogenic events leading to neurodegeneration in PD, notably mitophagy and protein degradation pathways (*2*). Here, we examined how disruption of the trafficking and delivery of CCVs by a mutation in the *DNAJC6* gene results in dopaminergic dysfunction and parkinsonism. By studying the loss of function mutations in Auxilin, we have also gained novel insight into its physiological activity and describe a novel role for Auxilin in the uncoating of Golgi-derived CCVs.

### A role for Auxilin in the uncoating of CCVs at the synapse and the Golgi apparatus

Auxilin is the major neuronal CCV uncoating protein (*9*). It is recruited to CCVs through membrane lipid interaction via its PTEN domain (Figure S11A) (*37*–*39*). In addition, Auxilin interacts with the clathrin lattice via clathrin binding boxes and with HSC70 via its J-domain (Figure S11A) (*12*, *38*–*40*). The subsequent stimulation of the ATPase activity of HSC70 by Auxilin then acts to drive the uncoating of the clathrin coat (*12*–*14*, *41*-*43*). In addition to these key interactions, Auxilin has previously been found to interact with the plasma membrane-resident clathrin adaptor protein AP2 (Figure S11A) (*16*, *17*, *24*, *26*), but not with the Golgi-resident AP1 (*16*). Here, in addition to AP2, we identified GGA2, a Golgi-resident clathrin adaptor protein, and AP180, a clathrin adaptor protein for synaptic vesicle recycling, as a novel *bona fide* Auxilin interactors (Figure 3, Figure S11A), thus describing for the first time a role for Auxilin in the uncoating of TGN-derived CCVs and a direct role in the recycling of SVs. Indeed, Auxilin was found to co-localize with the TGN and we observed an increase in CCVs both at the synapse and the Golgi apparatus in neurons with an endogenous missense mutation in Auxilin (Figures 5, 6). In contrast, we find that the ubiquitously expressed homologue of Auxilin, GAK, interacts with AP1 and AP2, but not the GGA proteins (Figure S7) (*16*). These results show that Auxilin and GAK interact with different Golgi-resident clathrin adaptors and suggest that the two homologues are only partially redundant functionally.

### Auxilin-associated mutations are loss of function mutations

Multiple recessive mutations in Auxilin have been associated with parkinsonism (Figure S11A). Two splice-site mutations (c.801-2A>G, T741=) are predicted to result in overall decreased expression of Auxilin and are thus hypomorphic (*4*, *5*). In addition, three C-terminally truncating mutations (Q791X, Q846X, R256X) completely lack the J-domain, required for interaction with HSC70, thus pointing to a partial loss of function mechanism (*6*–*8*). However, the mechanism of action of the R927G point mutation in the J-domain of Auxilin is less clear (*5*). We engineered a novel mouse model carrying an R857G substitution in Auxilin, homologous to the R927G mutation, and found that the R857G Auxilin allele is hypomorphic during early development, as demonstrated by decreased Auxilin protein levels in the brain of neonatal mice (Figure 1 D, E). Structural modeling of Auxilin indicated that the R927G mutation resides within the coiled-coil J-domain, thereby disrupting hydrogen bonds required for alpha-helical formation (Figure 4 B, C). In addition, all tested nonsense (Q791X, Q846X) and missense (R927G) Auxilin mutations were found to disrupt or weaken the interaction with its chaperone HSC70 and with clathrin (Figure 4, S7). These interactions are crucial for the correct positioning of HSC70 in close proximity to a subset of critical interactions within the clathrin lattice and for the stimulation of HSC70 ATPase activity for the uncoating reaction (*12*). It is thus conceivable that impaired clathrin interaction as well as disruption of the J-domain would impair CCV uncoating in Auxilin mutation carriers. Indeed, the number of CCVs was found to be drastically increased in the brain of R857G Auxilin mice (Figure 5), with an increase of coated structures both at the synapse and the Golgi apparatus (Figure 5, 6). These findings suggest a stalling in the clathrin uncoating reaction, with subsequent accumulation of CCVs in the brain of Auxilin mutation carriers and impaired delivery and recycling of intracellular vesicles.

### Neuropathological lesions underlie PD-like neurological phenotypes in R857G Auxilin mice

We observed age-dependent movement phenotypes in R857G Auxilin mice, including bradykinesia and gait disturbances, as well as seizures (Figure 2, Video S1), similar to those observed in patients. These symptoms occurred in the absence of gross neurodegeneration, but were accompanied by pathological changes in the brain, including intracellular lipofuscin-like accumulations in the nigrostriatal pathways of R857G Auxilin mice (Figure 7). Impaired clathrin uncoating of TGN-derived CCVs results in impaired delivery of its cargo, including proteins and lipids, to their intracellular destination compartments (Figure S11B). TGN-derived CCVs are particularly important for the delivery of hydrolases to the lysosomes (*44*). Inefficient delivery of those hydrolases would therefore decrease the neuronal degradative capacity and further aggravate the accumulation of intracellular cargo (Figure S11B), as has previously been reported for Auxilin depleted cells (*45*–*50*).

Clathrin-mediated trafficking also plays a major role in the recycling of synaptic vesicles (*51*). Impaired clathrin uncoating of synaptic CCVs results in inefficient recycling of SVs, indicated by the decreased number of pre-synaptic SVs as observed in the brain of R857G Auxilin mice (Figure 5, Figure S11B). Taken together, impaired clathrin-mediated trafficking in dopaminergic neurons, both at the synapse and the Golgi apparatus, may underlie the parkinsonism phenotypes, including progressive bradykinesia and gait disturbances, observed in R857G Auxilin mice (Figure S11B). In addition, impaired uncoating of CCVs in other brain areas might contribute to the epileptic seizures.

### Clathrin-mediated trafficking and Parkinson’s disease

In addition to Auxilin mutations, loss of function mutations in clathrin uncoating protein Synaptojanin 1 have also been associated with early onset PD. Patients with recessive mutations in the neuronal phosphatase synaptojanin 1, required for the shedding of clathrin adaptor proteins during CCV uncoating, present with similar phenotypes as described in Auxilin mutation carriers, including motor impairments and seizures (*52*). These findings underscore an important role for clathrin coat dynamics in early onset PD and broaden the spectrum of pathways and cellular processes linked to dopaminergic dysfunction in the mammalian brain, further emphasizing the importance of intracellular vesicle trafficking.

In this study, we describe a novel role for Auxilin in the uncoating of TGN-derived clathrin vesicles. Impaired lysosomal clearance and post-Golgi trafficking have previously been associated with PD pathogenesis, as multiple Mendelian genes including *LRRK2* and *VPS35* play prominent roles in vesicular trafficking between the Golgi apparatus and endosomes (*22*, *53*). In addition, multiple PD risk factors, including *GBA* and *CTSB*, are lysosomal hydrolases and require clathrin-mediated trafficking for their correct delivery to lysosomes (*20*, *21*).

At a time when there is an urgent unmet need for disease modifying therapies for Parkinson’s disease, our results further diversify the range of potential targets to be investigated and exploited. Response to L-DOPA, the first-line treatment in PD, is either absent or limited due to severe sideeffects in Auxilin mutation carriers (*4*–*8*). Further dissection of clathrin-dependent pathways in neurons is therefore of particular interest to find novel potential therapeutic targets. The murine model with endogenous PD-associated Auxilin mutation described herein provides a valuable platform to carry out such investigations, as well as to screen for potential disease modifying drugs for Parkinson’s.

## Material and methods

### R857G Auxilin mice

For the generation of the R857G Auxilin mice on a C57BL/6 background, CRISPR sgRNA (AAGTGAAGAAGGTGTACAGG) and oligonucleotides (GGAGACCAAATGGAAACCCG TGGGCATGGCGGATCTGGTGACGCCGGAGCAAGTGAAGAAGGTGTACGGCCGCGCTGTG CTAGTGGTGC ACCCTGACAAGGTGGGTAGCACCTGCCCTGTCGTAGACTTGCCCGGTCCCTGTTTCAGTGTTC) for CRISPR editing were designed using the web-based Benchling software (https://benchling.com). sgRNAs were selected based on their proximity to the PAM sequence and based on maximal on-target and minimal off-target effects (score system as described in (*54*)). Mouse mating pairs were set up on the day before micro-injection. Fertilized eggs were harvested and microinjected with Cas9 mRNA (12.5 ng/μl), sgRNA (4 ng/μl) and donor oligonucleotides (100 ng/μl). Zygotes were cultured overnight in M16 medium at 37°C and 2-cell stage embryos were implanted into oviducts of pseudo-pregnant surrogate mothers. Two male mice born to the foster mothers with successful homozygous gene editing were bred with C57BL/6J mice to establish the R857G Auxilin knockin mouse line. Mice were crossbred for at least 2 generations. The mice were given access to food and water *ad libitum* and housed in a facility with 12 hour light/dark cycles. All experiments were conducted in strict accordance with the recommendations in the Guide for the Care and Use of Laboratory Animals of the National Institutes of Health and approved by the Animal Care and Use Committees of the US National Institute on Aging.

### Behavioral analysis

All behavioural experiments were performed during the light cycle of the mice and all animals were handled for 2 minutes, 3 days prior to testing. The longitudinal cohort consisted of 8 WT and 8 R857G Auxilin mice that were age-matched, with 4 male and 4 female mice per genotype. Animals were subjected to behavioural tests at 2, 6, 12 and 18 months of age. *Beam walk* Mice were placed on an elevated narrow square beam of 100 cm in length with an enclosed dark platform at the end of the beam. Mice were trained on a beam of 12 mm in width for 3 consecutive trials on 3 consecutive days. On testing day, time to traverse a 12 mm or 6 mm beam was measured for 2consecutive trials.

*Rotarod* Mice were trained on a rotating rod for 5 minutes at a constant speed of 4 rpm. Starting the next day, mice were tested for three consecutive days on an accelerating rod from 4-40 rpm over 5 minutes. Latency to fall was determined 3 times for each mouse at 20 minute intervals and the average latency to fall was measured for each mouse per day.

*Pole test* Mice were placed head-upward on the top of a wooden dowel (1 cm diameter, 0.5 m height) and recorded by video as they descended. One pre-trial was performed followed by two test trials. Time to descend to the floor of the cage as well as time to turn head-downward was measured and averaged for the two test trials. Maximal score was assigned for mice that did not turn or climb from the pole for time to turn and time to descend, respectively.

*Grip strength* Mouse grip strength was measured using a digital grip strength gauge. The apparatus was connected to a wire grid of 8 by 8 cm. The mice were lifted by the tail to allow them to grasp the grid with their forelimbs. Mice were pulled backward gently by the tail until the grid was released. The peak full force in grams exerted by the mouse before losing grip was recorded. The mean of 5 consecutive trials was recorded for each mouse.

*Open field* Mice were allowed to habituate in the testing room under red light for at least an hour. Mice were then placed in a Flex field photobeam activity system with 25.4 x 47 cm dimensions consisting of 4 x 8 photobeams for 30 minutes. Activity was tracked by photobeam breaks in real time. Total activity count was measured as the arithmetic count of the total number of beam breaks, fine movement count as the number of single beam breaks and subtraction of the fine movement counts from total movement counts resulted in the ambulatory event count. Rearing counts indicated the arithmetic count of all beam breaks registered by a second level of photobeams. Path length was calculated based on the coordinates of the beam breaks. Activity in the center was calculated by breaks of photobeams 2-3 (out of 4 total horizontal beams) and photobeams 3-6 (out of 8 total vertical beams).

*Amplitude of movement* Mice were placed in an SR-Lab startle response system for 15 minutes, in a non-restrictive plexiglass cylinder (3.2 cm diameter) resting on the sensor platform within a sound- and light-proof box. A piezo-electric accelerometer was attached to the base of the sensor platform, thus converting mouse displacement and acceleration into a voltage measurement, which was digitized by the SR-Lab software. Voltages were measured every ms throughout the entire test. For analysis, voltage measurements as a function of time were Fourier transformed to extract frequency information using the ‘seewave’ and ‘rgl’ package for R. *Elevated plus maze* Mice were allowed to habituate to the dimly lit (100 lutz) room for an hour. Mice were then placed in an elevated plus-shaped maze, with each arm of the maze 38 cm in length and 10 cm in width. Two arms of the maze opposite to each other were enclosed with 15 cm high walls. The mice were placed in the center of the maze facing a closed arm and were allowed to explore the maze for 10 minutes. The number of arm entries, time spent in each arm and percentage of entries into the open arms was scored.

*Spontaneous alternation* Mice were placed in a symmetrical Y-maze consisting of 3 arms, each 40 cm long, 8 cm wide and enclosed by plexiglass walls that were 12 cm high. Mice were placed in the center of the maze and were allowed to explore all 3 arms of the maze freely for 8 minutes. Spontaneous alternation was defined as consecutive entries in 3 different arms divided by the number of possible alternations.

*Forced alternation* The forced alternation task was conducted in the same Y-maze as described above. The forced alternation task consisted of a 5 minute sample trial and a 5 minute retrieval trial, with a 90 minute inter-trial interval. During the sample trial, the mice were placed in the start arm and were allowed to explore 2 arms of the Y-maze, whilst the third arm was blocked. During the retrieval, this block was removed and the mouse was placed in the start arm and allowed to freely explore all 3 arms of the Y-maze. Forced alternation was scored as the percentage of mice in the retrieval trial entering the arm that was blocked during the sample trial first. In addition, time spent in the novel arm was measured.

### RNAscope

RNAscope was performed as described in (Wang et al., 2012). Probes were designed for DNAJC6 (target region 235-1177 of NM 001164583.1), GAK (target region 395-1305 of NM 153569.2) and the DA neuronal marker TH (target region 483-1603 of NM 009377.1).

### Mouse brain CCV isolation

A protocol for isolation of clathrin-coated vesicles with adapted from (*56*). Briefly, brains from 20 wildtype and 20 R927G DNAJC6 knockin mice between 2-4months old were homogenized using 7mL glass Dounce homogenizers in ice cold buffer A (100mM MES p.H. 6.5, 1mM EGTA, 0.5mM MgCl_2_, 200mM PMSF, and 1mg/mL pepstatin-A). Samples were centrifuged for 20 minutes at 20,000g to pellet debris (P1) using a SW 32Ti rotor. The remaining supernatant (S1) was centrifuged at 55,000g for 1hr. P2 was resuspended in buffer A and homogenized briefly before mixing equal parts buffer A with Ficoll and sucrose such that the final concentration was 6.25% of each. P2 was then centrifuged at 40,000g for 40min. The resulting supernatant was diluted 5 times in buffer A and respun at 100,000g for 1hr. P4 was resuspended in 6mL buffer A and homogenized briefly before centrifugation at 20,000g for 20min. Lastly, the supernatant containing CCVs and SVs were layered onto buffer A that was prepared with D_2_O and 8% sucrose and spun at 25,600rpm for 2hr. P6 was resuspended in 200μl of buffer A supplemented with 1X Cell Signaling Lysis buffer before using in Western blot analyses.

### Neuronal FM 1 43 dye uptake

Primary neurons from cortex were prepared from postnatal day 0 pups as described previously (*55*). Primary neurons were cultured until week 4 to reach a more mature state in supplemented BME as described above. FM1–43FX stryl dye (ThermoScientific) was solubilized in HBSS at 37°C to a stock concentration of 1 mg/mL. FM1–43 dye uptake was performed as described previously(*35*). In brief, cells were washed briefly with HBSS and then incubated at 37°C with HBSS media containing Ca^2+^ for 5 min. Media was aspirated and replaced with depolarizing solution (HBSS with Ca^2+^ containing 60 mM KCl^+^ along with 5 μg/mL FM 1–43FX stryl dye) for 1 min. Depolarizing solution was replaced with HBSS with Ca^2+^ containing 5 μg/mL of FM 1–43FX stryl dye for an additional 10 min during the recovery phase. Following incubation, cells were washed three times with 1 ml HBSS. Cells were fixed either right after or 1h after using 4% PFA and mounted prior to imaging as described above. A minimum of 4 pups per genotype were used to culture cortical neurons. Neurons were seeded at 1.5 × 106 cells per well across 4 coverslips per genotype.

### Transfection and immunofluorescence of primary neurons

Primary were plated onto coverslips precoated with poly-D-lysine (Neuvitro) at 0.5×10^6^ cells/wells as described. Primary cortical neurons transfected using Lipofectamine 2000 (Invitrogen), according to manufacturer’s instructions. 2 μg plasmid was transfected for a 12 mm coverslip in a single well of a 24-well plate of primary neurons cultured 7 DIV. Culturing media was replaced 4 hours after transfection and immunocytochemistry was performed 40 hours after transfection. Cells were fixed for 20 minutes in PBS containing 4% paraformaldehyde and 120 mM sucrose, followed by permeabilization for 15 minutes with 0.2% Triton diluted in PBS. Cells were blocked for 30 minutes with 3% FBS in PBS. Next, neurons were incubated at RT for 1 hour with primary antibodies diluted in PBS containing 1% FBS (CHC (Abcam, ab21679), GM130 (Abcam, ab169276), GLG1 (ThermoFisher Scientific, PA5-26838), TGN38 (Bio-Rad, ab10552)). Cells were washed 3 times with PBS. Cells were subsequently incubated with Alexa Fluor secondary antibodies (ThermoFisher Scientific) diluted in PBS buffer containing 1% FBS for 30 minutes, followed by 3 washes with PBS-CM buffer. All secondary antibodies were donkey host and used at 1:500 dilution. Coverslips were mounted on microscope slides using ProLong gold Antifade Mountant (ThermoFisher Scientific) and dried overnight at RT in the dark.

Immunohistochemistry was performed on the brains of 12 month old mice, transcardially perfused with saline. Brains were fixed in 4% PFA for 48 hour and subsequently transferred to a 30% sucrose solution. Fixed brains were cut into coronal 30 μm sections and stored in antifreeze solution (0.5 M phosphate buffer, 30% glycerol, 30% ethylene glycol) at −20°C until further processing. Brain sections were transferred to 24-well plate and washed from antifreeze solution with PBS twice for 10 min. Sections that were stained against DAT were subjected to antigen retrieval prior to immunostaining. Section were placed into Citric buffer (10mM sodium citrate, 0.05% Tween 20, pH 6.0) for 30 min at 80°C and were rinsed again afterwards with PBS buffer. All sections were then incubated in PBS containing 10% NDS, 1% BSA and 0.3% Triton for 30 minutes. Following blocking, sections were incubated primary antibodies (DAT (Abcam, ab111468), TH (Pel-freeze Biologicals, P40101-150), VMAT (ImmunoStar, 20042)) in antibody solution (1% NDS, 1% BSA and 0.3% Triton in PBS) overnight at 4°C. The next day, sections were rinsed three times with PBS for 10 min and incubated with AlexaFluor labeled secondary antibody (1:500, Invitrogen, Donkey host) in antibody solution for 1 hour. For neutral lipid staining, sections were incubated in 20 μg/ml BODIPY493/503 (Invitrogen). Afterwards, sections were washed three times with PBS for 10 min, mounted on glass slides and mounted using Prolong Gold Antifade mounting media (Invitrogen).

### Electron microscopy

Striatal brain slices were collected from 10 month old mice. Mice were transcardially perfused with saline for 2 minutes, followed by perfusion with fixation buffer for 5 minutes (2% formaldehyde, 2% glutaraldehyde in 150 mM sodium-cacodylate, buffered at pH 7.4). Brains were isolated and postfixed for 8 hours in fixation buffer. Next, brains were rinsed overnight in 150 mM sodium-cacodylate buffer without fixatives. The following day, 200 μm thick coronal brain sections were sliced using a vibratome. Striatal sections around the anterior commissure level were submitted for conventional transmission EM imaging. Specimens were rinsed in cacodylate buffer, postfixed with 1% OsO4 in the same buffer on ice, en bloc stained with 1% uranyl acetate, dehydrated in an ethanol series and embedded in EMbed 812 resin (Electron Microscopy Sciences). Thin sections were cut, stained with uranyl acetate and lead citrate, and viewed with a JEM-1200EX (JEOL) transmission electron microscope (accelerating voltage 80 keV) equipped with an AMT 6 megapixel digital camera (Advanced Microscopy Techniques).

### Confocal laser scanning microscopy with Airyscan detection

Airyscan imaging was performed in enhanced-resolution mode on a Zeiss LSM 880 Airyscan microscope equipped with a 63X, 1.4 NA objective. Raw data were processed using Airyscan processing in ‘auto strength’ mode with Zen Black software version 2.3.

### SILAC-based proteomics

HEK293FT cells were metabolically labeled in DMEM supplemented with dialyzed 10% FBS, supplemented with 12C6 L-Lysine-2HCl (Lys-0) and 12C6 L-Arginine-HCl (Arg-0) or 13C6 L-Lysine-2HCl (Lys-8) and 13C6 L-Arginine-Hcl (Arg-10) for the metabolic incorporation of ‘light’ and ‘heavy’ stable isotopes respectively (ThermoFisher Scientific). HEK293FT cells were grown in light or heavy SILAC media for at least 10 doublings, allowing for higher than 98% efficient incorporation of the light or heavy stable isotopes. HEK293FT cells labelled with ‘light’ or ‘heavy’ SILAC isotopes were transfected with GFP or GFP-Auxilin, respectively, using Lipofectamine 2000 according to manufacturer’s instructions. Co-immunoprecipitations were performed as described below and the resulting triplicate samples were loaded on a polyacrylamide gel for electrophoresis and subsequently cut out from the gel and subjected to in-gel trypsin digestion, followed by liquid chromatography tandem mass spectrometry analysis (LC-MS/MS). The LC-MS/MS data were searched against the NCBI Human database and Mascot Distiller software was used to calculate the protein Light/Heavy ratios (O’Leary et al., 2016; Perkins et al., 1999). Interactors identified across all 3 replicates were considered for further bio-informatic filtering. Proteins were filtered to have a Mascot protein score higher than 50, an FDR-corrected p-value < 0.05 and at least 4-fold enrichment in GFP-Auxilin samples compared to GFP control. In addition, the CRAPome database was used to filter out common contaminants of affinity purification-mass spectrometry experiments(*25*).

Functional enrichment analysis was performed for the 50 most significantly differentially expressed genes using Gene Ontology (The Gene Ontology Consortium, 2019; The Gene Ontology Consortium et al., 2000) for gene Ontology terms ‘biological process’ and ‘cellular component’. Fischer exact test was performed for functional enrichment analysis with Bonferroni post hoc correction. An enrichment map was generated using the ‘EnrichmentMap’ Cytoscape plug-in.

### Co-immunoprecipitation

HEK293FT cells were transfected with GFP/GFP-Auxilin plasmid using Lipofectamine 2000, as per manufacturer’s instructions. Co-IPs were performed 24 hours after transfection. Cells were resuspended lysis buffer (20 mM Tris pH7.5, 10% glycerol, 1 mM EDTA, 150 mM NaCl, 0.3% Triton) supplemented with 1x protease inhibitor cocktail and 1x phosphatase inhibitor cocktail (Halt) and rotated at 4°C for 30 minutes. Protein lysates were cleared by centrifugation for 10’ at 4°C at 21,000 g). 1% input samples were prepared by dilution in 1x Laemmli sample buffer and boiling for 5 minutes at 95°C. For co-IPs, equal amounts of protein lysates were mixed with GFP-Trap agarose beads (Chromotek) and rotated at 4°C for one hour. Beads were subsequently washed 5 times with lysis buffer and boiled for 5 minutes at 95°C in 1X Laemmli sample buffer.

Immunoprecipitation of endogenous Auxilin from mouse brain (2-3 months old mice) was performed by coupling Auxilin antibody (Novus, NBP1-81507) to Dynabeads™ Protein G (catalog 10009D, Thermo Fischer Scientific). One half of a brain was homogenized with a dounce homogenizer in 2 mL lysis buffer containing protease inhibitors. Lysates were incubated on ice for 20 minutes and cleared by centrifugation 10 min at 4°C at 16,000 g. Lysates were subsequently mixed with Dynabeads and rotated at 4°C for 4 hours. Beads were then washed 4 times with lysis buffer containing 1% Triton. Samples were boiled 5 minutes at 95°C in 1X Laemmli sample buffer prior to WB analysis.

### Western blot

Brain hemispheres were homogenized in lysis buffer (20 mM Tris pH7.5, 10% glycerol, 1 mM EDTA, 150 mM NaCl, 1x protease inhibitor cocktail (Halt), 1x phosphatase inhibitor cocktail (Halt)) with 1% Triton using glass homogenizers and samples were lysed on ice for 20’. Protein lysates were subsequently cleared (10’ centrifugation at 4°C at 21 kg). Protein lysates were diluted to obtain final sample concentration of 30 μg per sample and boiled in 1x Laemmli Sample buffer (Bio-Rad). Protein samples were loaded on pre-cast 4-20% TGX polyacrylamide gels (Criterion, Bio-Rad). Electrophoresis was performed in 1x pre-mixed electrophoresis buffer (10 mM Tris, 10 mM Tricine, 0.01% SDS, pH 8.3, diluted with water) using the Criterion Vertical Electrophoresis Cell (Bio-Rad). Following gel electrophoresis, samples were transferred to Western blots were transferred to 0.45 μm pore-size nitrocellulose membranes (Bio-Rad) using the Trans-Blot Turbo Transfer System (Bio-Rad). Blocking of membranes was performed in a 1:1 solution of phosphate buffered saline (PBS) and Odyssey Blocking Buffer (Li-Cor). After blocking, membranes were incubated with primary antibodies diluted in antibody buffer (1:1 of Tris buffered saline (TBS) with 0.1% Tween and Odyss Blocking Buffer (Li-Cor)) overnight with gentle agitation at 4°C. Primary antibodies used for WB analysis were Auxilin (Novus Biologicals, NBP1-81507), GAK (Gift from Dr. Lois Greene), CHC (Abcam, ab21679), AP2 (Abcam, ab2730), AP1 (B & D Biosciences, 610385), GGA1 (Abcam, ab10551), GGA2 (Abcam, ab10552), GGA3 (B & D Biosciences, 612311), GFP (Sigma, ab290), GST (Sigma, 27-4577-01), Flag (Sigma, F1804). Following primary antibody incubation, membranes were washed 3 times for 5 minutes in TBS-0.1% Tween. Membranes were then incubated with fluorescent secondary antibodies (IRDye, Li-Cor) diluted 1:15000 in antibody buffer for 1 hour at room temperature (RT) under gentle agitation. Secondary antibody incubation was followed by 3 washes of 5’ each in TBS-0.1% Tween. Western blots were imaged using the Odyssey CLx system (Li-Cor) and quantified using Image Studio software.

### Structural modeling

Structural models were generated using PyMOL (Version 2.0, Schrodinger, LLC). Modeling of the interaction between the Clathrin triskelion and the J-domain of Auxilin was performed utilizing the structure from Bos Taurus (PDB:1XI5) emphasizing the pathogenic mutation and interaction domain (*12*). Representational modeling of potential interaction between GGA2 and Auxilin was performed using the previously solved structures (PDB: 3N0A-Auxilin and the S. Cerevisiae GGA2 PDB: 3MNM) and based on predicted interaction sites (Figure 6.7) (*16*, *37*, *57*, *58*). I-TASSER was utilized to predict the structure of human Auxilin and the strongest match was selected. The generated model was based on the Auxilin structure from Bos taurus (PDB: 3N0A) (*37*, *59*–*61*). Heterodimeric complex (Hsc70 and Auxilin) was modelled in PyMol overlaying the J-domain of the generated model with the Hsc70-J-domain structure available in the Protein Data Bank (PDB: 2QWO) (Figure 6.13) (*62*).

## Acknowledgments

This research was supported in part by the Intramural Research Program of the NIH, National Institute on Aging.

DAR was supported by a University of Reading Graduate School Doctoral award. PAL is supported by MRC programme grant MR/N026004/1 and the Aligning Science Across Parkinson’s research network (ASAP 0478).

This work was supported, in part, by the NIMH IRP Rodent Behavioral Core (ZIC MH002952).

## Author Contributions

DAR, PAL and MRC conceptualized the research. CL generated the CRISPR KI mice. Structural models were generated by NS. YL performed mass-spectrometry analysis. CKEB performed electron microscopy. LBP generated the SILAC HEK cells and performed mouse brain IPs. NL performed perfusions, IHC and FM-dye experiment. JK performed CCV isolation experiment. SSA performed RNAscope experiment. All other experiments were performed by DAR. MCM, JDH and JD contributed to the analysis of behavioral assays. CDW, DCG and JSB contributed to enhanced-resolution light microscopy. DCG and LS contributed to the analysis of EM experiments. AB, RK and AK contributed primary neuronal culture experiments. All experiments were analyzed by DAR. DAR, PAL and MRC wrote the manuscript.

