## Supplementary Figures for "Mutations in Auxilin cause parkinsonism via impaired clathrin-mediated trafficking at the Golgi apparatus and synapse"

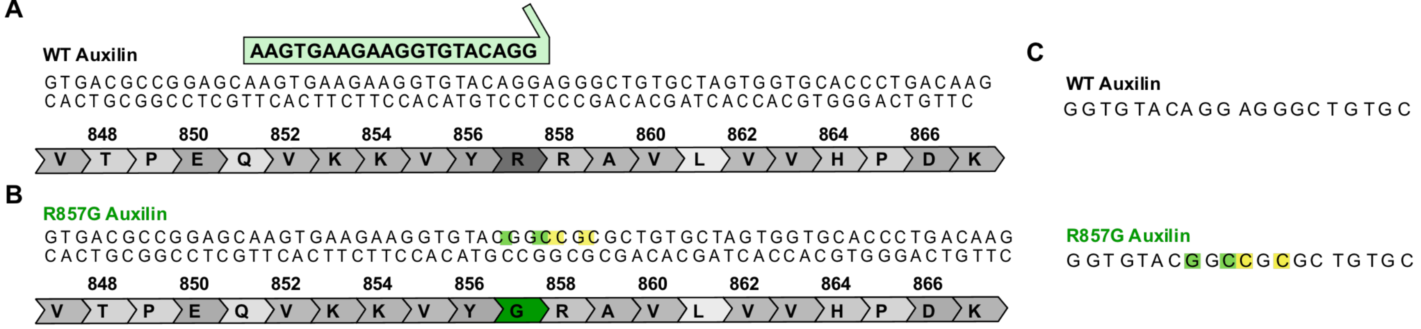


**Figure S1: CRISPR design of R857G Auxilin mouse** A, B Design of the R857G Auxilin mutant mouse at the level of amino acids and nucleotides with indication of the gRNA sequence utilized for CRISPR editing. Silent mutations are indicated in yellow, missense mutations are indicated in green. C Sanger sequencing confirmed correct knocking of the R857G Auxilin mutation into the mouse genome.

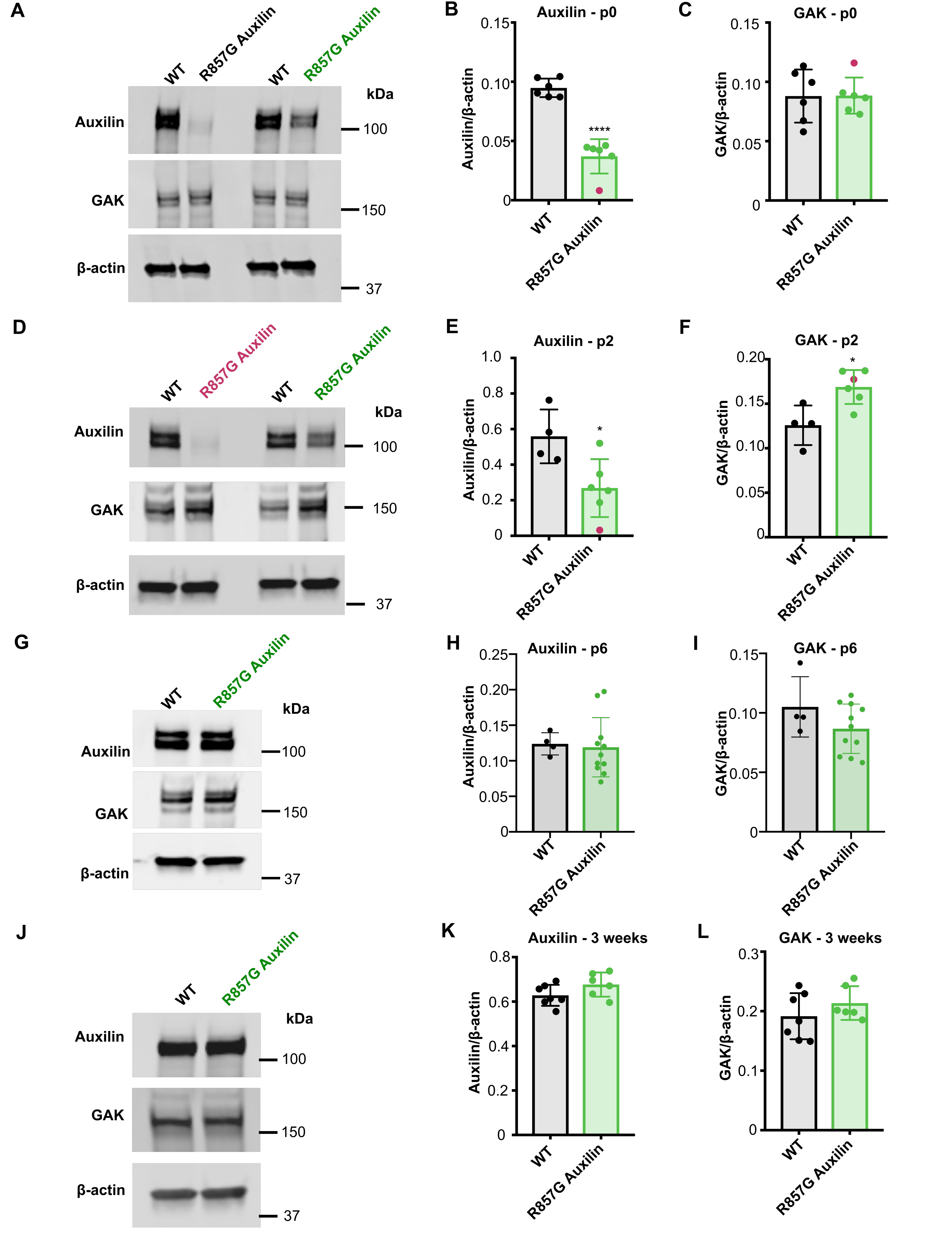


**Figure S2: Alterations in Auxilin and GAK protein levels in the brain of R857G Auxilin mice** A, D, G, J WB of Auxilin and GAK in brain lysates of P0, P2, P6 and 3 week old mice, respectively. N = 4-6 per genotype. B, C, E, F, H, I, K, L Quantification of normalized Auxilin and GAK levels in the brain lysates o WT and R857G Auxilin mice. Welch's t-tests were performed and p-values indicated when significant, with * $p < 0.05$ and **** $p < 0.0001$.

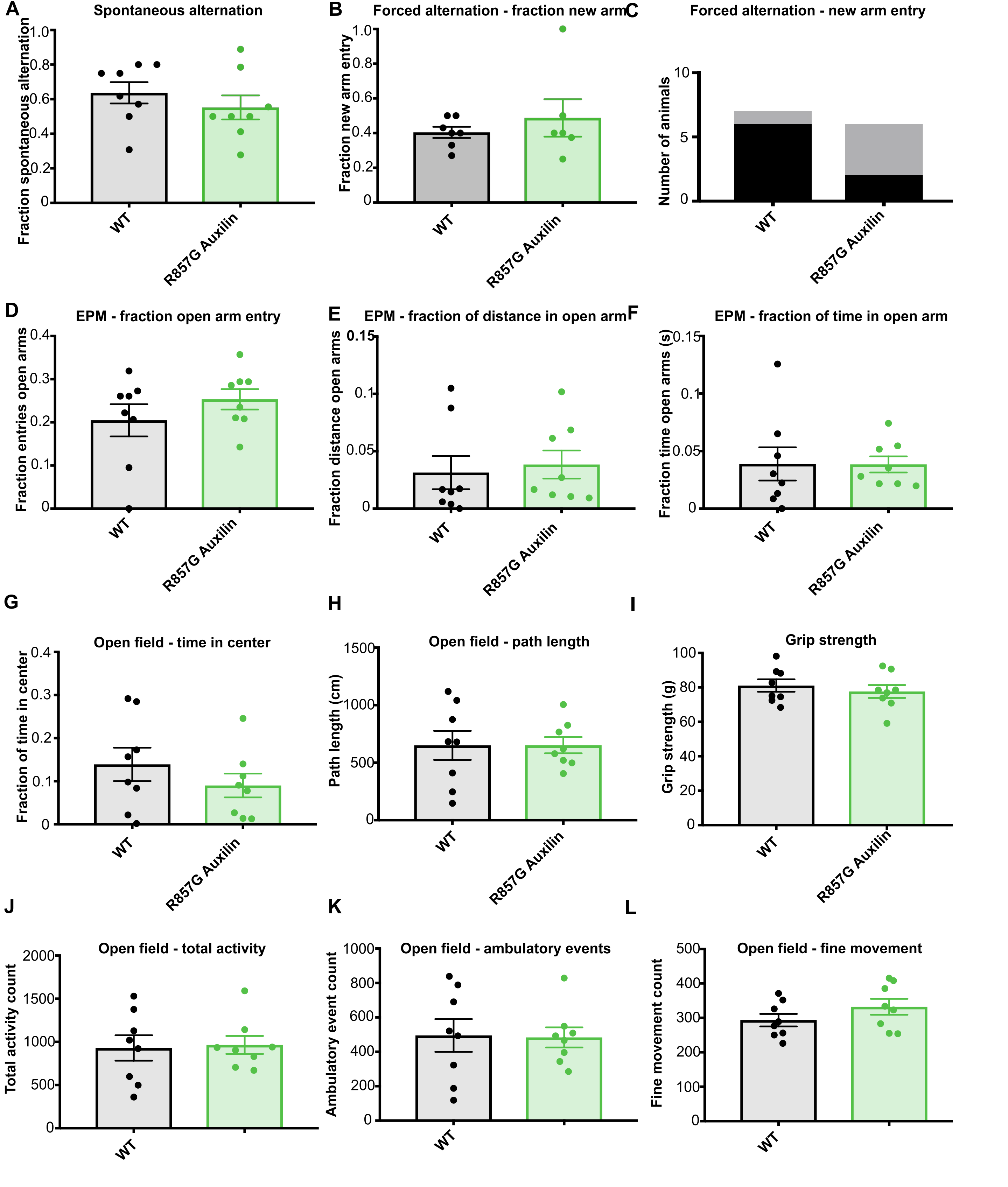


**Figure S3: Anxiety, memory and locomotor phenotypes in 1 year old R857G Auxilin mice** A Y-maze spontaneous alternation test indicating the fraction of spontaneous alternations. B Y-maze forced alternation test indicating the fraction of entries the new arm. C Y-maze forced alternation test indicating the number of entries into the new arm upon first arm entry. D, E, F Fraction of open arm entries, fraction of distance travel in open arm and fraction of time spent in open arm of the elevated plus maze, respectively. G Time spent in the center of the open field test. H Total path length in the open field test. I Average forelimb strength measured during 5 trials of the grip strength test. J, K, L total activity, ambulatory activity and fine movement counts, respectively, of WT and R857G Auxilin mice in the open field test.

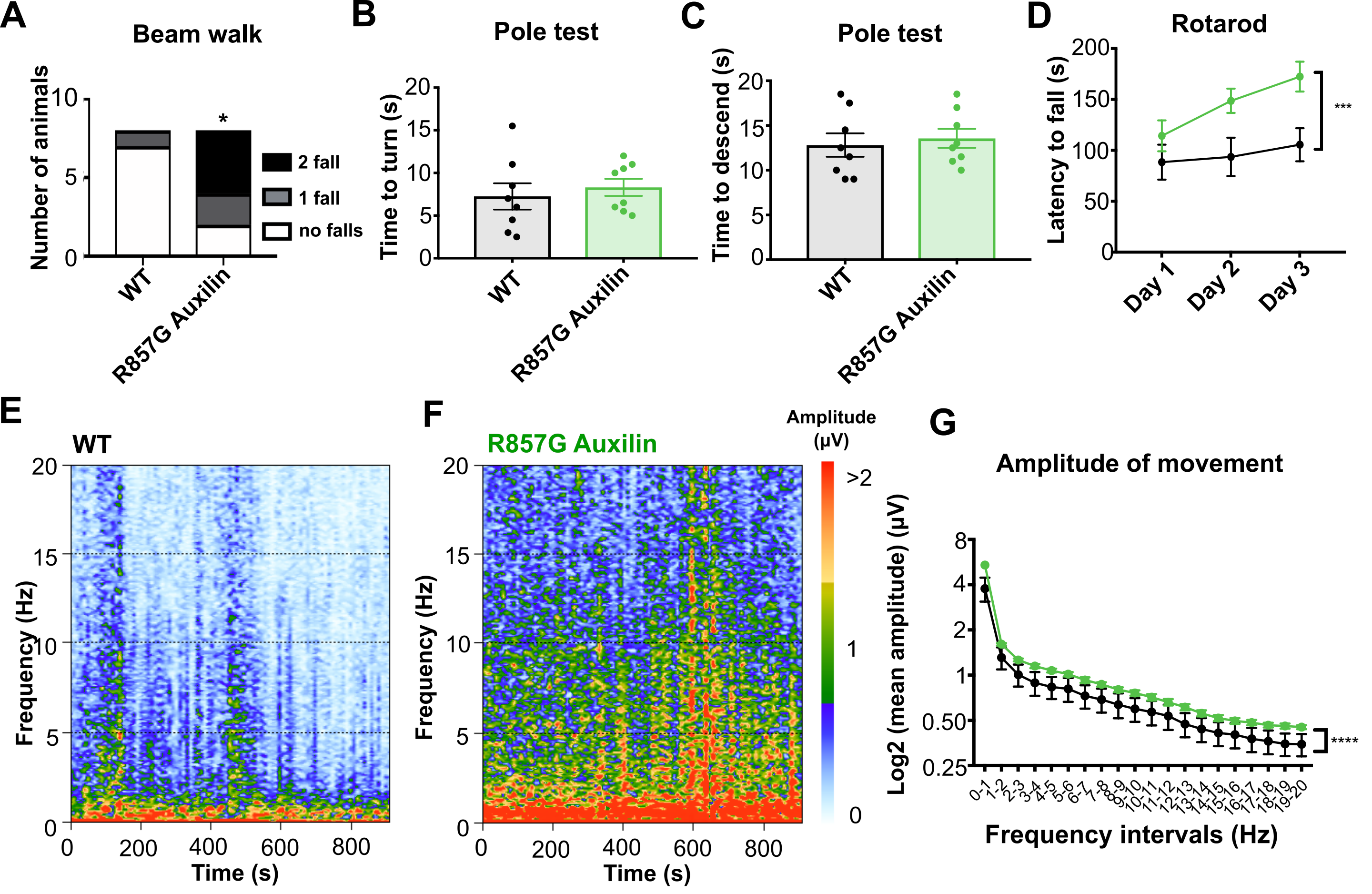


**Figure S4: Motor impairments in 6 month old R857G Auxilin mice** A Beam walk analysis indicating the number of falls when traversing a 6 mm beam during 2 trials. Chi-square tests were performed, * indicates $p < 0.05$. B, C Performance during the pole test as analyzed by time to turn and time to descent. D Latency to fall during the rotarod test during 3 consecutive days. Two-way ANOVA was performed with *** $p < 0.001$. E, F Analysis of the amplitude of movement in a startle chamber. Representative spectrograms of WT and R857G Auxilin mice are shown, displaying the amplitude (indicated by color) of movement during 15 minutes over a frequency range of 0-20 Hz. G Two-way ANOVA was performed, with **** $p < 0.0001$.

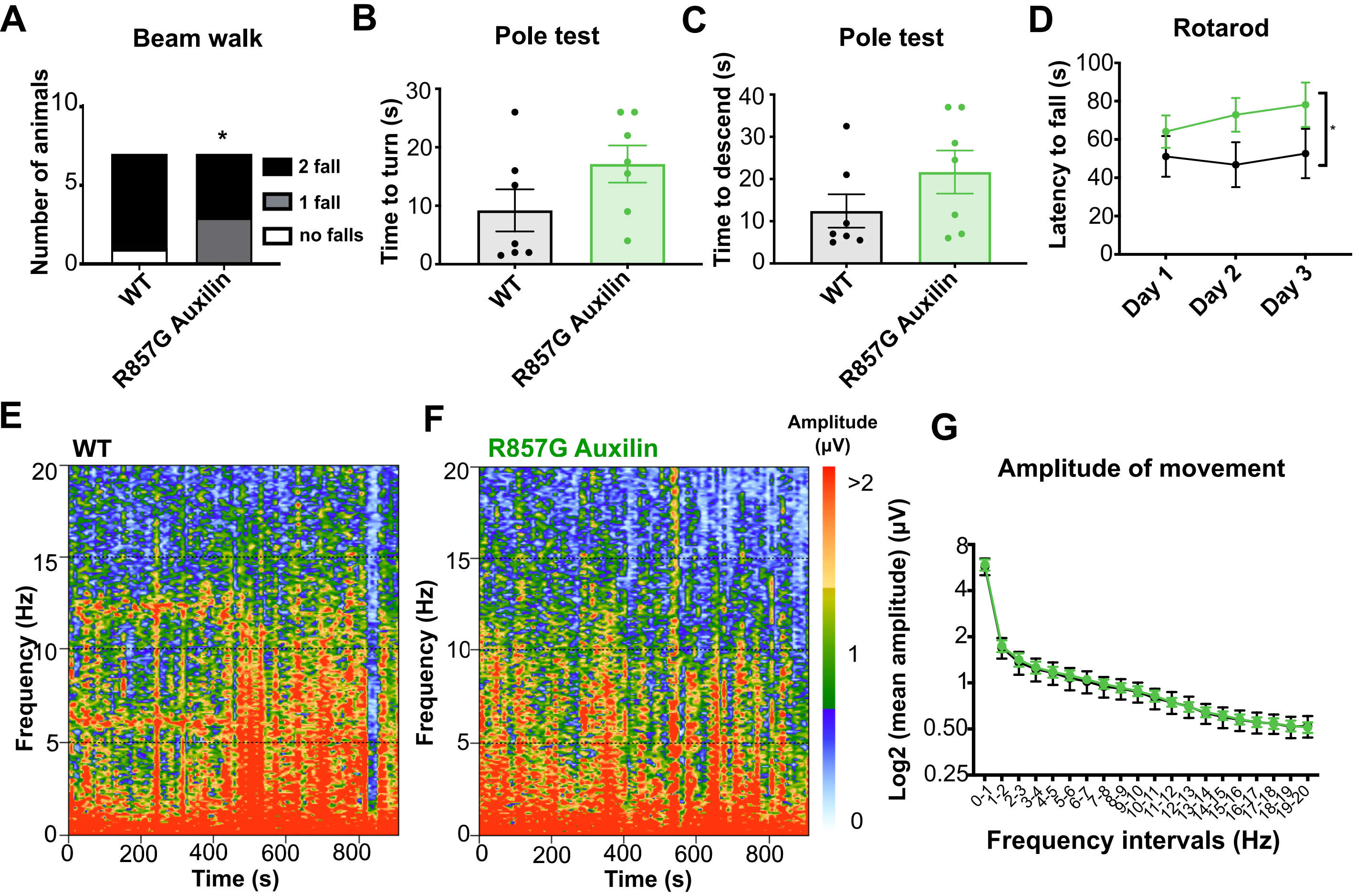


**Figure S5: Motor impairments in 18 month old R857G Auxilin mice** A Beam walk analysis indicating the number of falls when traversing a 6 mm beam during 2 trials. Chi-square tests were performed, * indicates $p < 0.05$. B, C Performance during the pole test as analyzed by time to turn and time to descent. D Latency to fall during the rotarod test during 3 consecutive days. E, F Analysis of the amplitude of movement in a startle chamber. Representative spectrograms of WT and R857G Auxilin mice are shown, displaying the amplitude (indicated by color) of movement during 15 minutes over a frequency range of 0-20 Hz. G Average amplitude of movement over a frequency range from 0-20Hz.

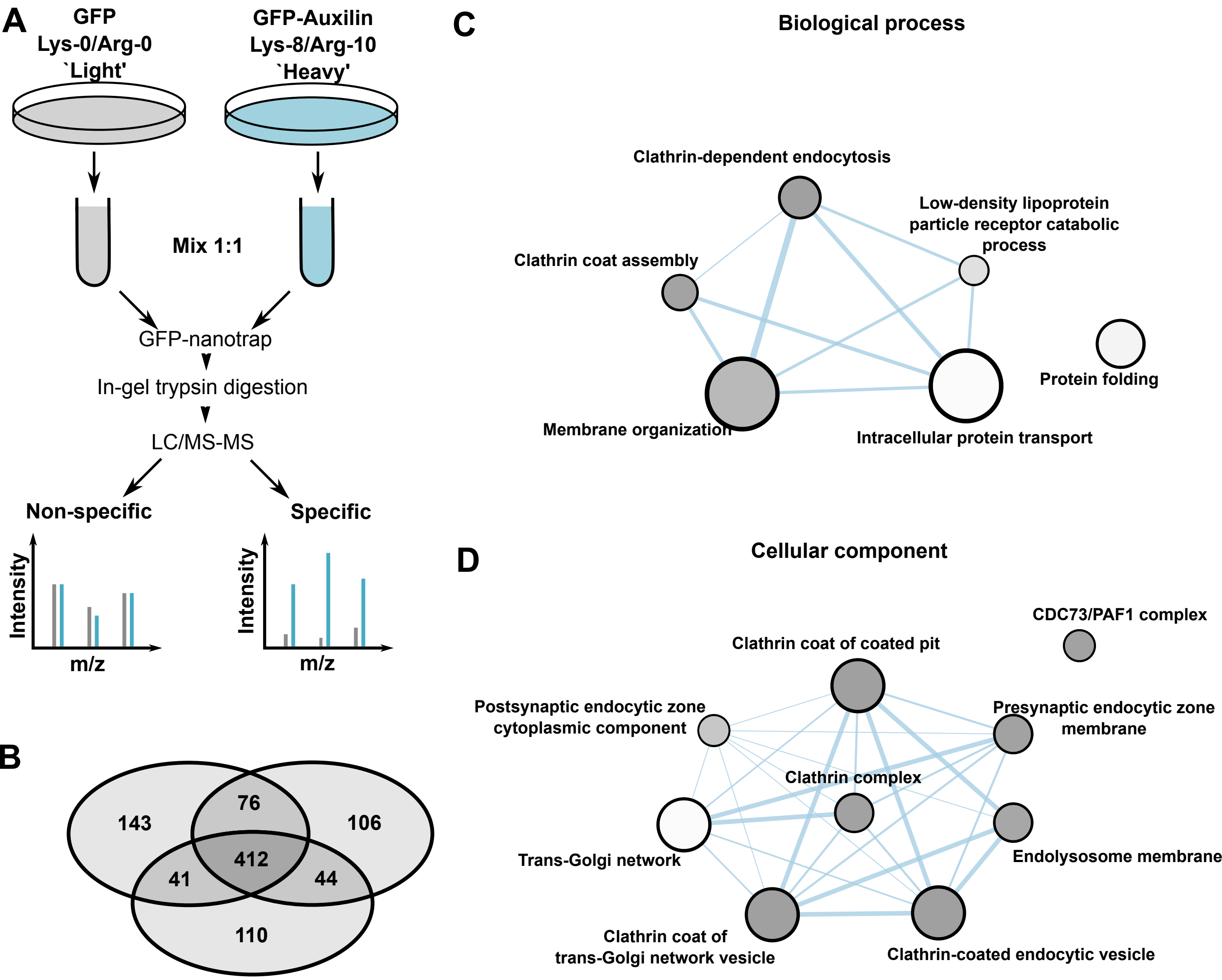


**Figure S6: SILAC set-up and gene ontology analysis of the Auxilin interactome** A Design of SILAC-based AP-MS approach to identify the interactome of Auxilin. HEK293FT cells were labelled with 'light' or 'heavy' amino acid epitopes and transfected with GFP or GFP-Auxilin, respectively. GF nanotrap co-IP was performed and precipitates were mixed 1:1, followed by in-gel trypsin digestion, LC-MS/ MS and subsequent protein identification. B Venn diagram of the number of proteins identified across 3 technical replicates of the SILAC experiment. C, D Gene ontology enrichment analysis of the top candidates of Auxilin interactors for biological process and cellular component. Node size corresponds to the number of proteins within each protein set and edge size to the number of overlapping proteins between connected nodes (larger is more proteins). Node color corresponds to the p-value of the Fisher exact test with Bonferroni correction for multiple testing (darker is lower p-value).

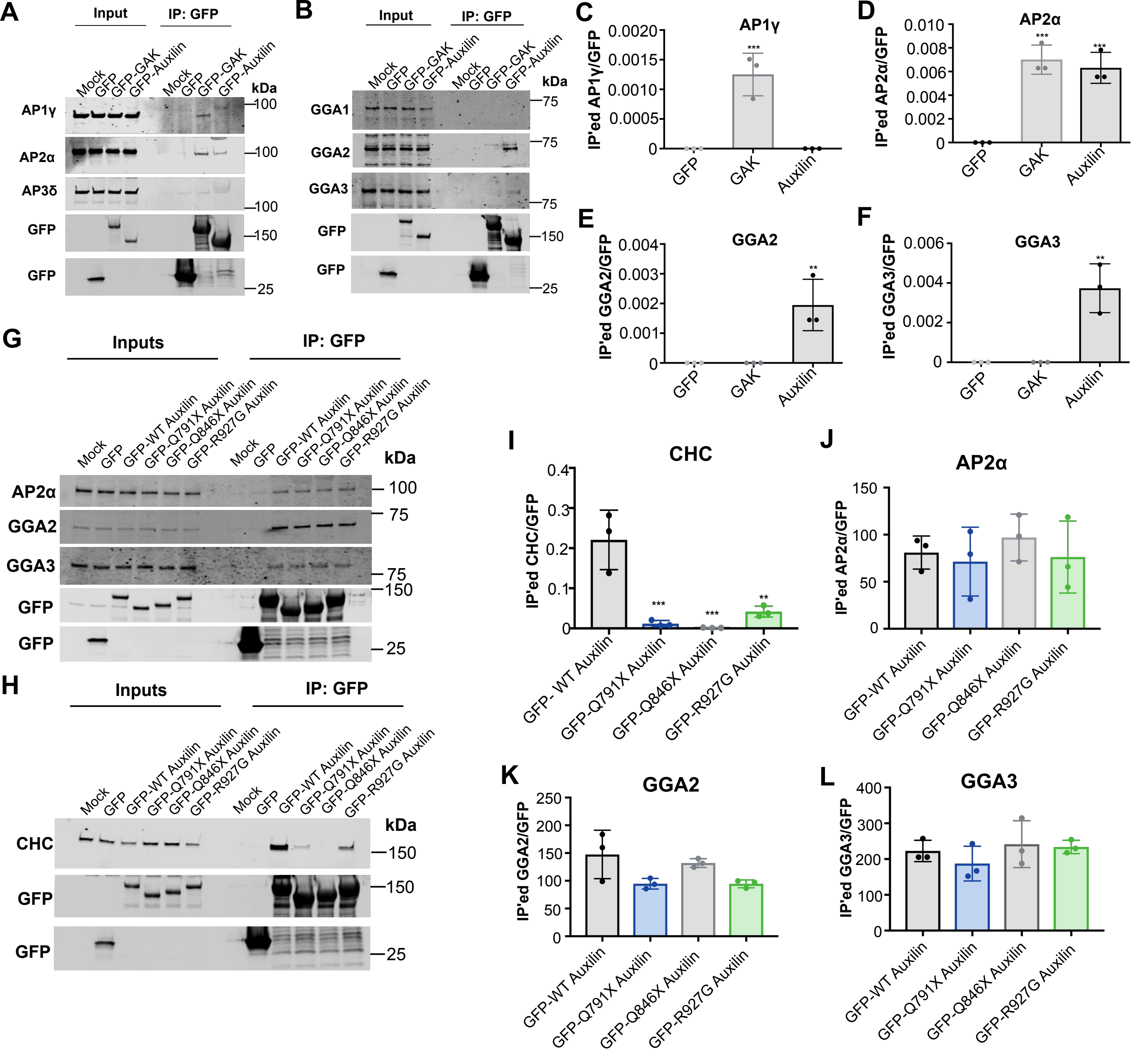


Figure S7: A, B GFP-GAK or GFP-Auxilin were transiently expressed in HEK293FT cells and WB analysis was performed of GFP-nanotrap experiments for interaction with endogenous AP and GGA proteins. Images are representative of n = 3 technical replicates. C-F Quantification of AP1, AP2, GGA2, GGA3 interaction with GAK/Auxilin compared to GFP negative control and normalized to bait. One-way ANOVA was performed with Tukey's post hoc test, p-values indicate *** $p < 0.001$, ** $p < 0.01$. G,H GFP or WT, Q791X, Q846X, R927G Auxilin were transiently expressed in HEK293FT cells and WB analysis was performed on GFP-nanotrap experiments for interaction with endogenous CHC, AP2, GGA2 and GGA3. Images are representative of n = 3 technical replicates. I-L Quantification of CHC, AP2A2, GGA2 and GGA3 interaction of mutant Auxilin compared to WT and normalized to bait. One-way ANOVA was performed with Tukey's post hoc test, p-values indicate *** $p < 0.001$, ** $p < 0.01$.

| | | | |
|---|---|---|---|
| **A** | **GAK** | **956** | AAADP**FGPL**LPSSGNNSQPCSNPDL**FGEF**LNSD |
| | **Auxilin** | **639** | ATFDPFGAPSKPSGQ - - - - - - - -DLLGSFLNTSS |
| **B** | **GAK** | **625** | DGKAVIPLGVTVQGDVLIVIYHARSTLGGRLQAKMASMKMFQIQFHTGFVPRNATT |
| | **Auxilin** | **338** | DGKI**FIPL**NITVQGDVVVSMYHLRSTIGSRLQAKVTNTQIFQLQFHTG**FIPL**DTTV |
| **C** | **AAAA Auxilin** | **338** | DGKI**A**IP**A**NITVQGDVVVSMYHLRSTIGSRLQAKVTNTQIFQLQFHTG**A**IP**L**DTTV |

**Figure S8: Interaction motifs for γ-ear containing accessory proteins in GAK and Auxilin** A Binding motifs in GAK for AP1 binding are indicated in bold and are not conserved in Auxilin [15,53]. B Putative binding motifs in Auxilin for GGA2 are indicated in bold and are not conserved in GAK [53]. C Mutation of putative GGA2 binding motifs in Auxilin, with mutated residues indicated in red.

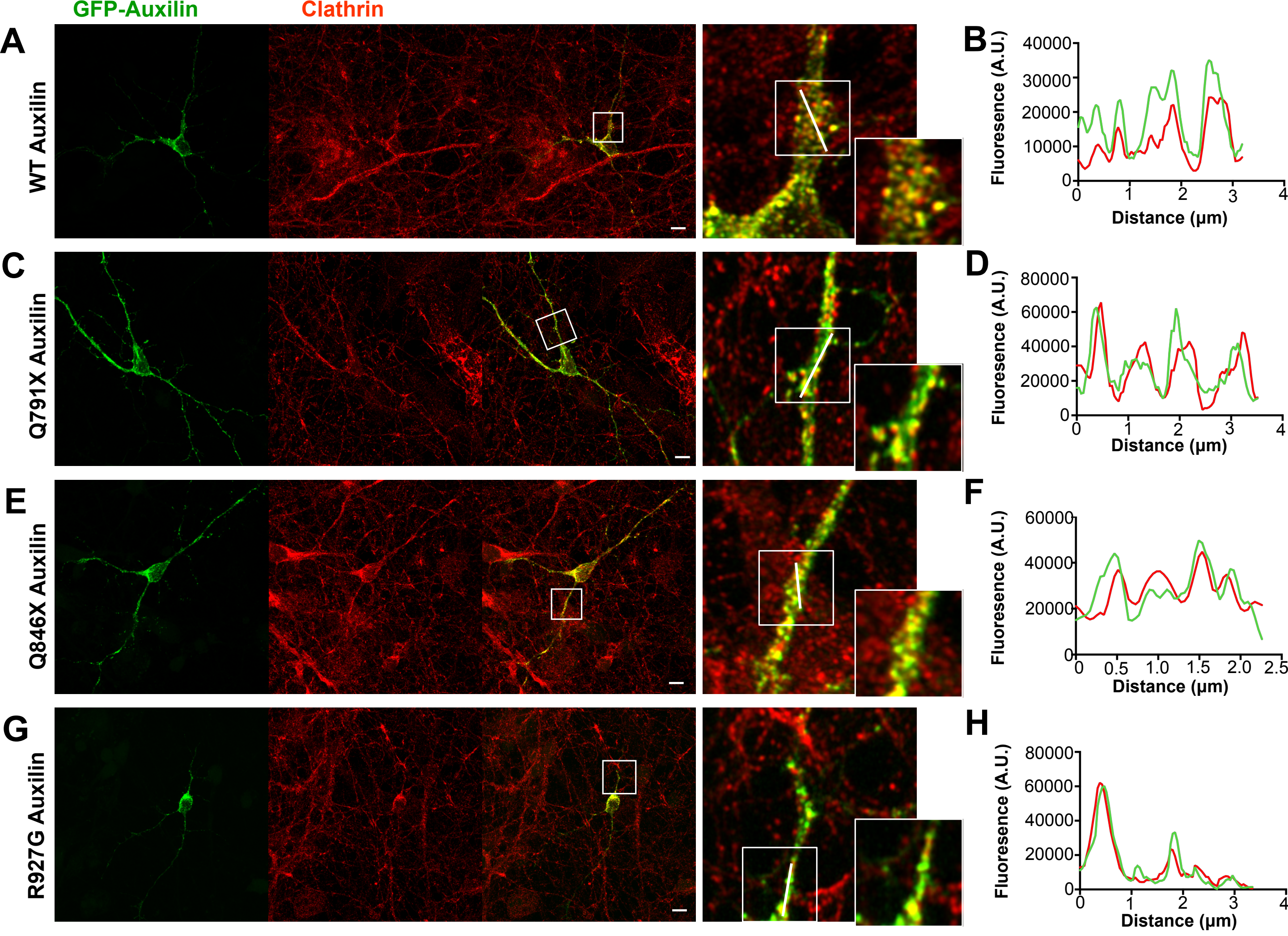


**Figure S9: Co-localization of Auxilin with clathrin** GFP-WT, Q791X, Q846X or R927G Auxilin (green) were transiently expressed in murine primary neurons and co-stained for endogenous clathrin heavy chain (red). Scale bar = 20 µm. The right panel shows the fluorescence intensity profiles in function of the distance indicated in the magnified merged image. Scale bar = 2 µm

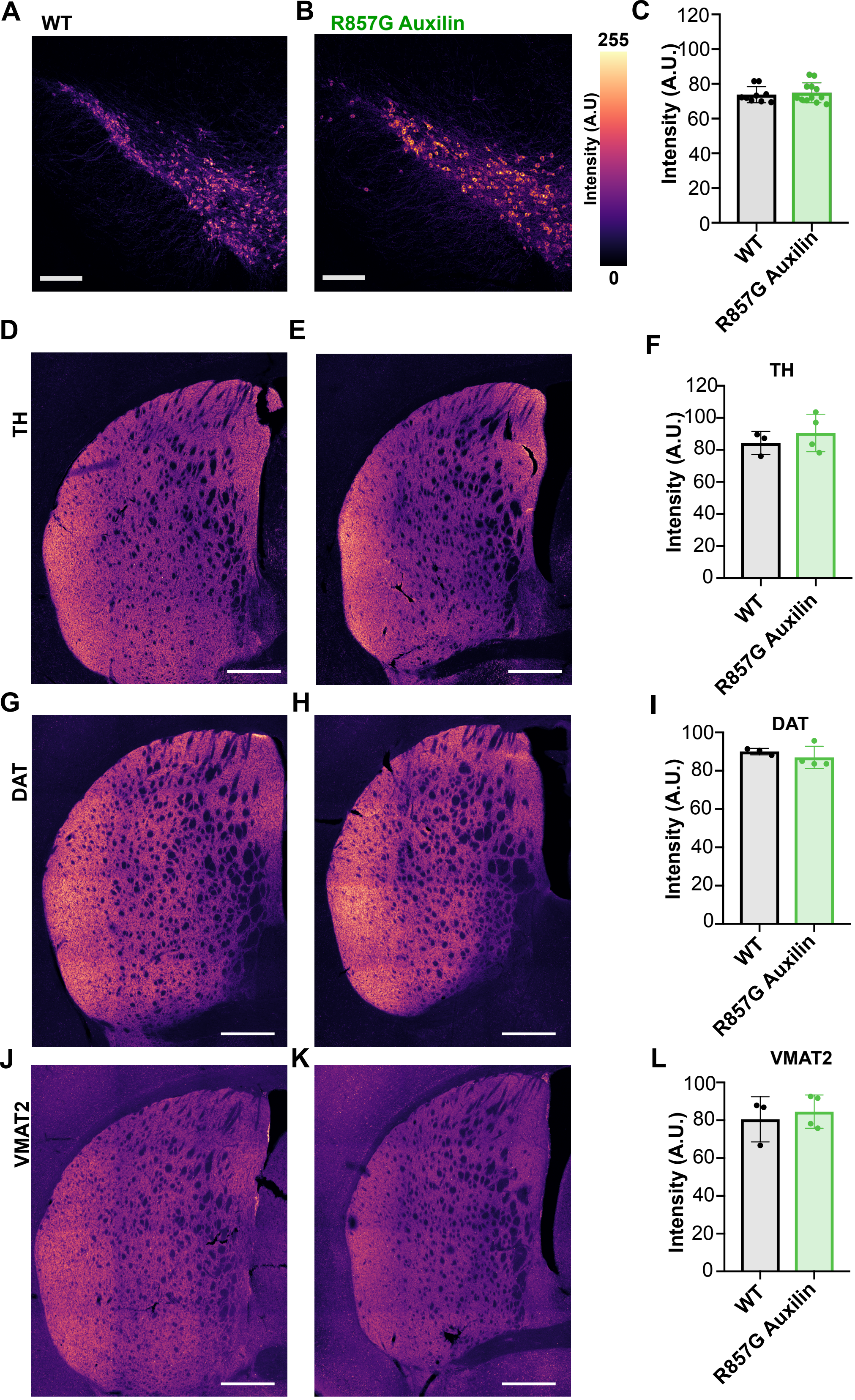


**Figure S10: No dopaminergic neurodegeneration in 12 month old R857G Auxilin mice** Staining for TH in the SN of WT and R857G Auxilin mice (A, B). Representative images are shown of 3 sections of n = 3 WT and n = 4 R857G Auxilin mice. Scale bar indicates 200 µm. Staining for TH (D, E), DAT (G, H) and VMAT2 (J,K) in the striatum of WT and R857G Auxilin mice. Representative images are shown of n = 3 and n = 4 WT and R857G Auxilin mice, respectively. Scale bar indicates 500 µm. Fluorescence intensity (F, I, L) was quantified and Welch's t-test was performed. No significant alterations were observed.

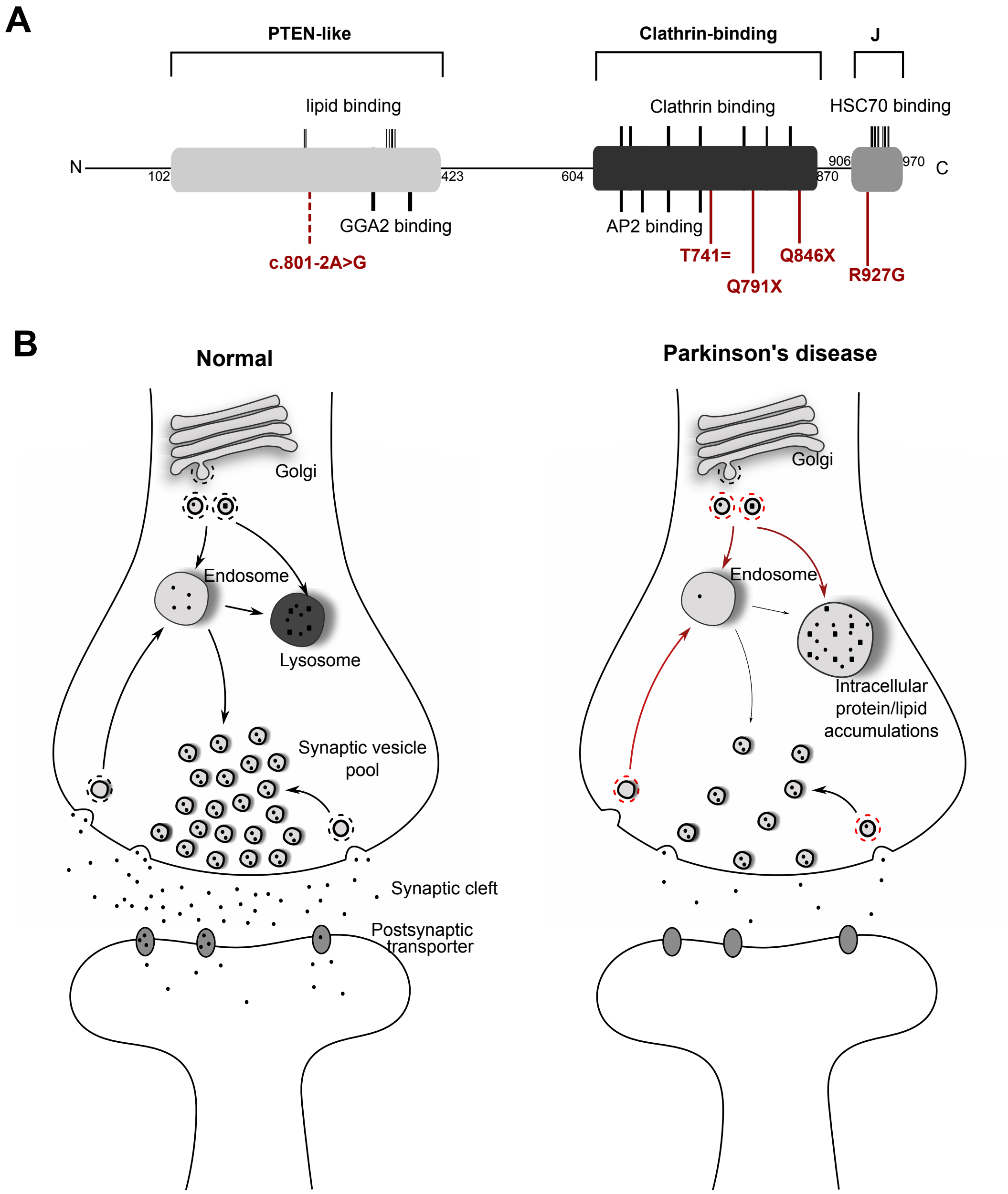


**Figure S11: A Overview of bona fide Auxilin interactors and interaction motifs. B Model for disease pathogenesis in Auxilin mutation carriers** A domain organization of Auxilin with indication of interaction motifs and residues involved with interaction for lipid, clathrin, HSC70, GGA2 and AP2 binding. Thickness of lines misrepresentative for number of residues involved with the interaction. Pathogenic mutations are shown in red, with solid line indicating coding variants and dotted line indicating exonic variant. B Model for PD pathogenesis in Auxilin mutation carriers Clathrin-mediated trafficking in neurons of normal population and PD patients. Impaired clathrin-mediated trafficking due to PD-associated Auxilin mutations would result in inefficient synaptic vesicle recycling, with subsequent decrease in number of vesicles in the pre-synaptic intracellular synaptic vesicle pool. In addition, inefficient clathrin-mediated trafficking from the TGN would result in impaired delivery of lysosomal hydrolases and therefore impair lysosomal maturation, resulting in an accumulation of intracellular cargo.
